# A requirement for Krüppel-Like Factor-4 in the maintenance of endothelial cell quiescence

**DOI:** 10.1101/2022.05.09.491221

**Authors:** Victoria Mastej, Cassondra Axen, Anita Wary, Richard D. Minshall, Kishore K. Wary

## Abstract

**Rationale and Goal:** Endothelial cells (ECs) are quiescent and critical for maintaining homeostatic functions of the mature vascular system, while disruption of quiescence is at the heart of endothelial to mesenchymal transition (EndMT) and tumor angiogenesis. Here, we addressed the hypothesis that KLF4 maintains the EC quiescence.

**Methods and Results:** In ECs, KLF4 bound to KLF2, and the KLF4-transctivation domain (TAD) interacted directly with KLF2. KLF4-depletion increased KLF2 expression, accompanied by phosphorylation of SMAD3, increased expression of alpha-smooth muscle actin (αSMA), VCAM-1, TGF-β1 and ACE2, but decreased VE-cadherin expression. In the absence of Klf4, Klf2 bound to the *Klf2*-promoter/enhancer region and autoregulated its own expression. Loss of EC-*Klf4* in *Rosa^mT/mG^::Klf4^fl/fl^::Cdh5^CreERT2^* engineered mice, increased Klf2 levels and these cells underwent EndMT.

**Conclusion:** In quiescent ECs, KLF2 and KLF4 partnered to regulate a combinatorial mechanism. The loss of KLF4 disrupted this combinatorial mechanism, thereby upregulating KLF2 as an adaptive response. However, increased KLF2 expression overdrives for the loss of KLF4, giving rise to an EndMT phenotype.

**Key Points:** Adult endothelial cells (ECs) are quiescent in that these cells are arrested at G_0_-phase of the cell cycle, but mechanisms of EC quiescence are not well understood.

The Krüppel-like factors (KLFs) -2 and -4 are transcriptional regulators, highly expressed in quiescent ECs, however, their roles in this process have not been addressed.

Elucidation of the mechanisms of KLF function in quiescent ECs should provide clues to the translational discoveries intended for the treatment of EC-dysfunction, such as endothelial to mesenchymal transition (EndMT) associated with several vascular diseases including tumor angiogenesis.

**Graphical Abstract:** 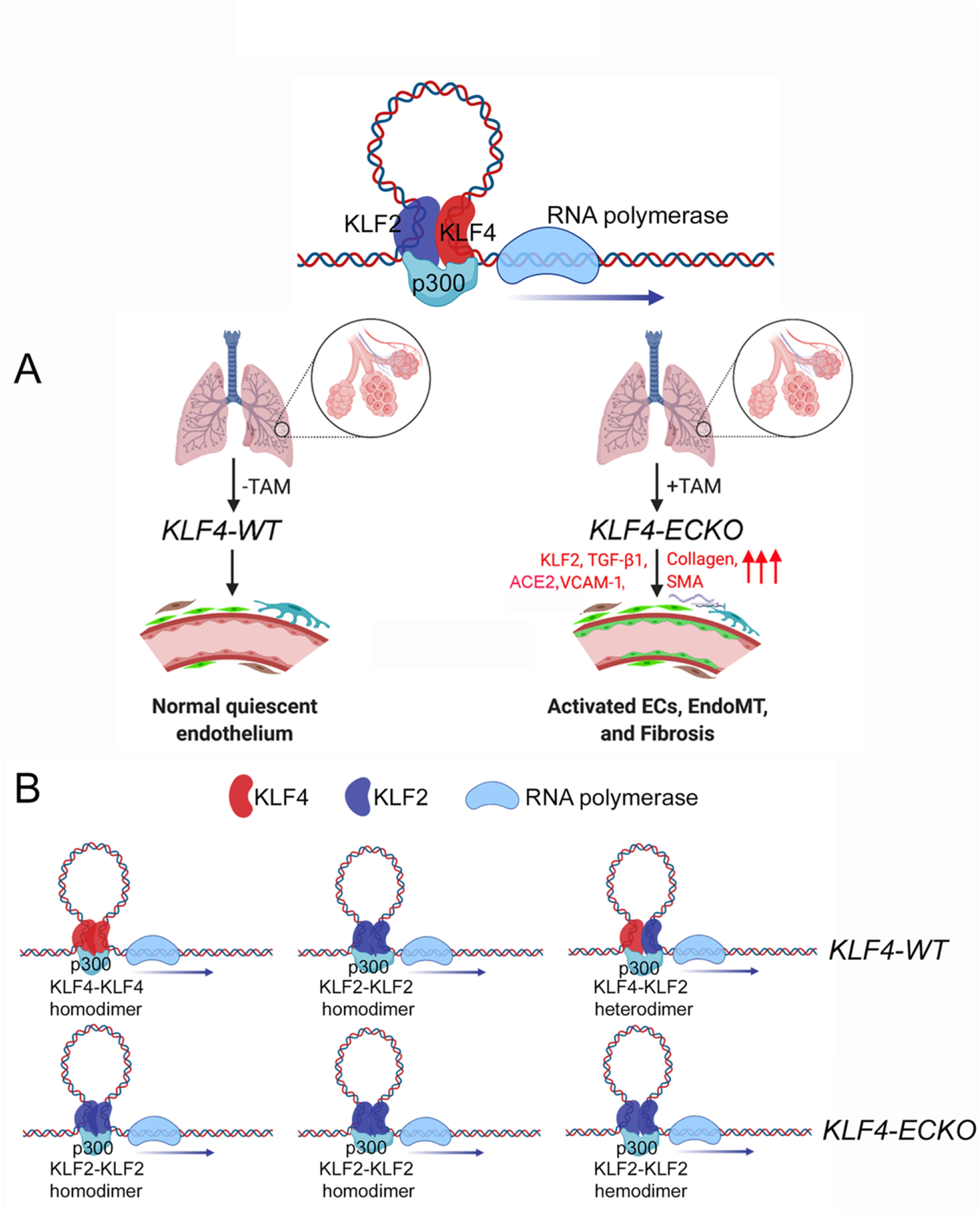

## Introduction

In the adult, ECs are highly specialized, heterogenous, and form a continuous monolayer via cell-cell and cell-matrix interactions (1–4). These interactions are the defining hallmarks of EC quiescence (4–8). EC quiescence is regarded as the default cellular state for a mature EC monolayer (1,4,5,7,8). Quiescent ECs critically regulate the anti-adhesive, anti-inflammatory, and anti-thrombotic behavior of quiescent blood vessels. Conversely, the disruption of EC quiescence in response to regenerative stimuli or injury from virus, bacteria, or angiogenic factors are likely to be at the heart of EC dysfunction, EndMT, systemic sclerosis (SSc, or scleroderma) including tumor angiogenesis (9–19). Importantly, EndMT is associated with the development of resistance to anti-cancer therapy, cardiotoxic effects of anti-cancer drugs; however, the molecular mechanisms of EndMT is not completely understood.

Hypoxia and tumor angiogenic factors, such as vascular endothelial Growth Factor (VEGF) can activate and transform quiescent ECs into proliferative stalk cells, this event propagates signaling that results in EndMT, produce *de novo* fibroblast and myofibroblast activation, extracellular matrix (ECM) deposition, and tissue fibrosis (15–19). Analogous to epithelial to mesenchymal transition (EMT) (20), EndMT is associated with the loss of VE-cadherin localized in adherens junctions, increased expression of mesenchymal transcription factors, loss of cellular polarity, and activation of TGF-β signaling frequently seen in melanoma and Kaposi sarcoma (21–24). Deletion of EC transcription factor, such as *Twist* and *Snail,* inhibited EndMT and ameliorated kidney fibrosis in mouse model (25). Silencing of epigenetic modifier JMJD2B in ECs reduced TGF-β2-induced expression of mesenchymal genes, limited the extent of EndMT (26). Thus, chronic inflammation and immune cell dysfunction are associated with EndMT (21–26). The loss of quiescent phenotype and the acquisition of EndMT are also associated with pulmonary arterial hypertension (4-6,27-29). Thus, chronic inflammation and sustained release of cytokines propagate signaling crosstalk, release of proteases, degradation of ECM components are considered hallmarks of EndMT and tumor microenvironment. However, the mechanisms of these processes are incompletely understood.

Transcription factors (TFs) KLF2 and KLF4 are highly expressed in quiescent ECs (30–36). To this end, KLF4 is known to inhibit cell cycle progression, while KLF2 inhibited VEGF-induced hyperpermeability. Importantly, KLF4 and KLF2 are expressed in differentiated endothelial and epithelial cell subpopulations (33,34,36–42). Both KLF2 and KLF4 contain a DNA binding domain composed of conserved C2H2 zinc fingers, a transactivation and repressor domain, and a nuclear localization signal (36,38,39,43). Additionally, KLF2 and KLF4 both share common regulatory mechanisms in that they recruit p300-CBP coactivator protein (44–46). Here, we addressed the hypothesis that EC-KLF4 and KLF2 maintain the quiescent state of ECs, while the loss of EC-KLF4 give rise to EndMT.

## Results

### 1. Expression of KLF4, KLF2, ACE2, and TGF-β1 associated with pulmonary microvessel remodeling in emphysema and COPD patients

KLF4 and KLF2 are abundantly expressed in normal adult lungs tissues (Fig. S1). The status of KLF2 and KLF4 expression in human chronic lung inflammatory disease have not been reported. To address this gap, lung tissue sections prepared from healthy control donors, emphysema and COPD patients were stained with anti-KLF2 or anti-KLF4 antibodies (Fig. S2A-D). Microscopic analyses of lung tissue sections showed thickening of alveolar structures in emphysema and COPD patients, compared with healthy vessels in donors (Fig. S2A-D). Quantification of KLF4 and KLF2 staining data in these tissue sections suggest that in normal lung, the level of KLF2 is relatively lower than KLF4 (Fig. S2E), in contrast, in emphysema and COPD tissue sections, the levels of KLF2 is higher than KLF4 (Fig. S2E). Total cell lysates prepared from a limited number of tissue samples, were subjected to Western blotting (WB) with antibodies against KLF4, KLF2, ACE2, TGF-β1, VE-cadherin and GAPDH. As VE-cadherin is exclusively expressed by mature ECs, we included VE-cadherin to demonstrate that presence of ECs in lung tissue samples we analyzed (Fig. S2F). We included TGF-β1, as previous studies have shown the ability of epithelial cell KLF4 to transcriptionally induce the expression of TGF-β1, thereby inducing epithelial to mesenchymal transition (EMT). Quantification of signal intensities of these proteins suggest that, the level of KLF4 is relatively higher in control lung tissues, while the expression of KLF2 increased and KLF4 decreased. Importantly, we detected increased ACE2 expression in lung disease tissue samples. As such, we also found evidence of EndMT in diseased lung tissue, as illustrated by co-alignment of anti-α-smooth muscle actin (αSMA) in CD31+ vascular structures (Fig. S3). Although, we surveyed limited number of tissue samples, together these data provided us the impetus to test the hypothesis that oscillation of KLF4 and KLF2 expression might be associated with EC-dysfunction and EndMT in the lung.

### 2. Oligomeric KLF4, KLF2, and p300 protein complexes in quiescent ECs

Most transcription factors form homo- or heterodimeric protein complexes. KLF2 and KLF4 have been shown to associate directly with p300, a nuclear protein that functions as a histone acetyltransferase (HAT) which regulates transcription via chromatin remodeling (44–46) but is not known to promote the formation of homo- or heterodimeric protein complexes. To test the hypothesis that KLF4 and KLF2 form multi-molecular complexes in human lung microvascular (hLMVECs), RIPA buffer solubilized nuclear lysates prepared from these cells were subjected to: a) Western Blot (WB) analysis and b) reciprocal coimmunoprecipitation experiment using anti-IgG (control), anti-KLF2, anti-KLF4, and anti-p300 antibodies, thereafter analyzed by WB immunoblotting with indicated antibodies. Equal amounts of proteins were boiled in native or reducing sample buffer and analyzed for occurrence of oligomeric polypeptide species (Fig. 1A-D). We detected presence of mono-, di-, and tetrameric anti-KLF4 and anti-KLF2 polypeptide species by native SDS-PAGE whereas in presence of strong reducing conditions (5% β-mercaptoethanol), only monomeric KLF4 and KLF2 were detected (Fig. 1A-D). Next, WB analysis was used to detect anti-p300 polypeptide immunoreactive species in anti-KLF2, anti-KLF4 and anti-p300 immunoprecipitated complexes (Fig. 1E). WB analysis with anti-KLF2 antibody showed the presence of ∼42kDa protein in anti-KLF4, anti-KLF2, and anti-p300 immunoprecipitates and anti-KLF4 antibody detected a ∼58kDa signal in anti-KLF4, anti-KLF2, and anti-p300 immunecomplexes; control IgG did not precipitate any of the above polypeptide species (Fig. 1E-G). To address equal loading across the lanes, total nuclear lysates (inputs) were analyzed by anti-KLF4 and anti-lamin-B1 antibodies (Fig. 1H&I). Together these data show that KLF2, KLF4, and p300 likely form mono-, di- or an oligomeric protein complexes in hLMVEC quiescent ECs (Fig. 1J).

**Figure 1.**
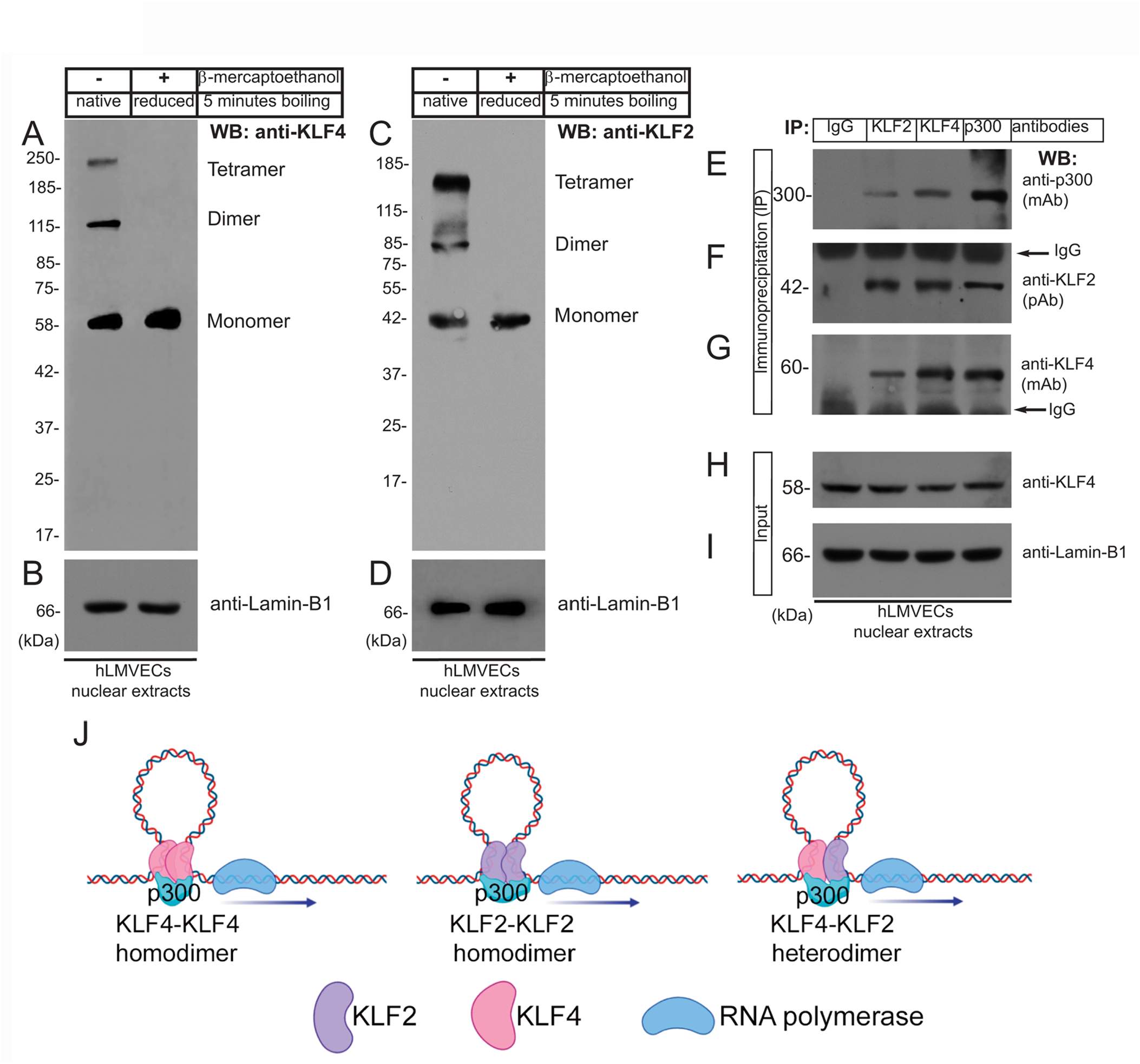
Oligomeric KLF4 and KLF2 protein complexes in quiescent ECs. **A&C**. Equal amounts of nuclear extracts prepared from human lung microvascular endothelial cells (hLMVECs) were boiled in native (with no β-mercaptoethanol) or reducing (with 5% β-mercaptoethanol) sample buffers, resolved by 9% SDS-PAGE, and analyzed for the occurrence of oligomeric protein complexes. The presence of mono-, di-, and tetrameric anti-KLF4 and anti-KLF2 polypeptide species in native SDS-PAGE were analyzed by indicated antibodies. **B&D**. Equal loading of nuclear extracts were analyzed by anti-Lamin-B1 antibody. **E-G.** Nuclear lysates prepared from hLMVECs were subjected to immunoprecipitation (IP) with IgG (control), anti-KLF2, anti-KLF4, and anti-p300 antibodies, and analyzed by WB with indicated antibodies. **H&I**. Nuclear lysates (input) were analyzed by WB with anti-KLF4 and anti-Lamin-B1 antibodies. Molecular weights are given in kiloDalton (kDa). This experiment was repeated at least three times. **J**. Models showing homodimer and heterodimer KLF2 and KLF4 protein complexes.

### 3. Transactivation domain (TAD) of KLF4 binds to KLF2

To address which segment is responsible for interaction of KLF4 with KLF2, we engineered a series of retroviral constructs encoding human KLF4-cDNA wild-type and deletion mutants, performed transfections, and conducted immunoprecipitation experiments using Flag-epitopes (-DYKDDDDK-, 1012 Da) that were fused in-frame to the C-terminus (Fig. 2A). The methods and efficiency of lentivirus vector pLNCX2-mediated transfection have been previously described by us (47–49). The strategy and timeline of transfection of pLNCX2 retroviral constructs are shown in Fig. 2B. hLMVECs at 50% confluence were used for this set of experiments. Nuclear extracts prepared from hLMVECs expressing a) pLNCX2 vector alone, b) human KLF4-WT cDNA, c) KLF4-Δ-86 construct (lacking 85 amino acid residues from the N-terminus), d) KLF4-Δ-96 construct (lacking 95 amino acid residues from the N-terminus), or e) KLF4-Δ-116 construct (lacking 115 amino acid residues from the N-terminus) and subjected to WB analyses showed the presence of corresponding polypeptide species (Fig. 2C). Equal loading was determined by WB analyses with anti-lamin-B1 antibody (Fig. 2D). Next, nuclear cell extracts that were immunoprecipitated with anti-Flag protein antibody were immunoblotted with anti-KLF4 antibody revealed bands for KLF4-WT (b), KLF4-Δ-86 (c), and KLF4-Δ-96 (d), while in control and KLF4-Δ-116 (e) samples, no band were detected (Fig. 2E). The non-specific signal at ∼52 kDa represents IgG (heavy chain). Stripping and re-probing the WB shown in Fig. 2E with anti-KLF2 Ab showed presence of KLF2 (Fig. 2F). Fig. 2G shows that an equal amount of nuclear extract was loaded in each lane (input). This data suggests that KLF4 likely binds to KLF2 via amino acid residues 97-117 present within the KLF4 transactivation domain (TAD).

**Figure 2.**
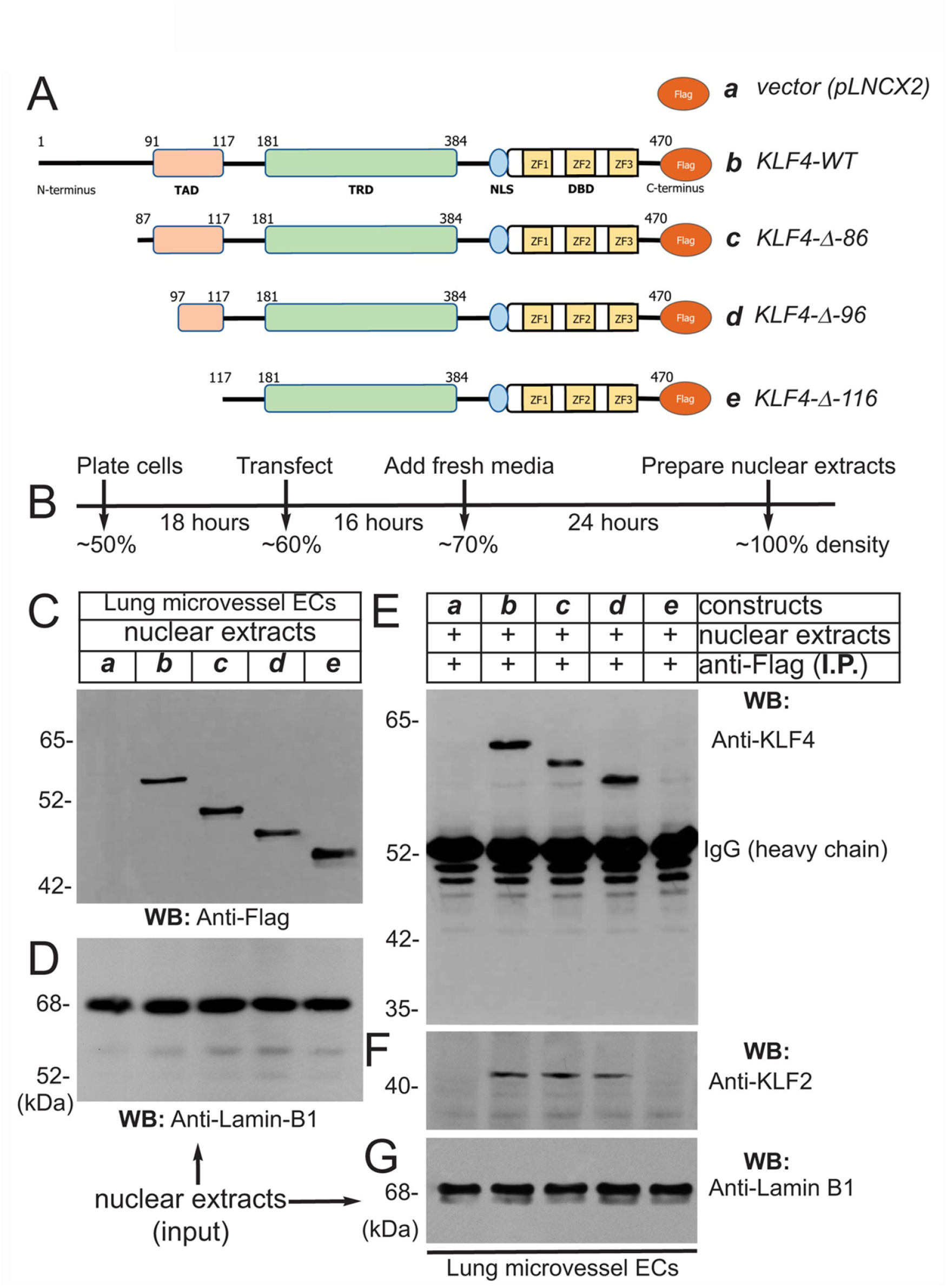
Transactivation domain (TAD) of KLF4 binds to KLF2. **A.** pLNCX2 retroviral vector encoding: a) vector alone; b) human *KLF4-wild-type (WT) cDNA* fused in-frame with Flag-epitope tag; c) *KLF4-Δ-86* construct (lacking 85 amino acid residues from the N-terminus) fused in-frame with Flag-epitope tag; d) *KLF4-Δ-96* construct (lacking 95 amino acid residues from the N-terminus) fused in-frame with Flag-epitope tag; e) *KLF4-Δ-116* construct (lacking 115 amino acid residues from the N-terminus) fused in-frame with Flag-epitope tag as shown. **B.** Strategy and timeline of human lung microvessel endothelial cell (hLMVECs) infection/transfection experiments. **C**. Efficiency of retrovirus mediated infection of hLMVECs was determined by WB with anti-Flag antibody. **D**. Equal amount of protein loading across the lanes were judged by anti-Lamin-B1 antibody. **E.** Equal amount of nuclear cell extracts were immunoprecipitated with an anti-Flag (2.0μg/30mg protein extract) monoclonal antibody (mAb) and analyzed by WB with anti-Flag mAb. **F**. The *KLF4-wild-type, KLF4-Δ-86,* and *KLF4-Δ-96* immuneprecipitated KLF2 polypeptide species, while *KLF4-Δ-116* did not. **G**. Input nuclear extracts showing equal levels Lamin-B1 proteins across the lanes. These experiments were repeated at least 3 times.

Next, to address if the interaction of KLF4 and KLF2 is direct or indirect, we carried-out GST pull-down experiments (Fig. 3). The amino acid sequence of human KLF4 (wild-type full-length) is shown in Fig. 3A. The GST-cDNA was fused in-frame to the C-terminus of the human *KLF4*-*cDNA* segment encoding amino acid sequence 91-116 (TAD), shown in blue (construct-b), and to region 181-200 (TRD), shown in magenta (construct-c). These cDNA constructs were used for preparation of GST-fusion proteins expressed in protease-deficient *E. coli* (BL2), thereafter purified by Sepharose-4B column chromatography (Fig. 3B&C). Next, nuclear extracts prepared from hLMVECs were used for GST-pull down experiments. The NC-membrane analyzed by WB showed the presence of expected anti-KLF2 antibody reactive ∼38 kDa polypeptide species (Fig. 3D). The amount of GST-fusion protein in each sample was determined by anti-GST antibody to insure equal loading (Fig. 3E). This data shows that KLF4-TAD protein interacts with KLF2, while KLF4-TRD segment did not. In addition, KLF4-TAD amino acid sequence (aa 91-117) was subjected to Iterative Threading ASSEmbly Refinement (I-TASSER) computational analysis. I-TASSER is a hierarchical method to protein structure prediction and structure-based function annotation (50,51). Accordingly, prediction revealed that amino acid residues 91-117 in KLF4 contain two discrete α-helical domains and two short stretches of coiled-coil regions (Fig. 3F-G). Further homology search identified 4E-BP1 that is structurally homologous to the KLF4 aa 91-117 secondary structure. 4E-BP1 is a translational repressor protein that binds to eukaryotic initiation factor 4E (eIF4E) and represses protein translation by inhibiting the ability of eIF4E to recruit 40S ribosomal subunits, for details please see computational analyses in supplement section (Fig. S4). Thus, we identified a new protein interacting domain within KLF4-TAD that has an intrinsic ability to form homodimer, heterodimer and oligomeric protein complexes.

**Figure 3.**
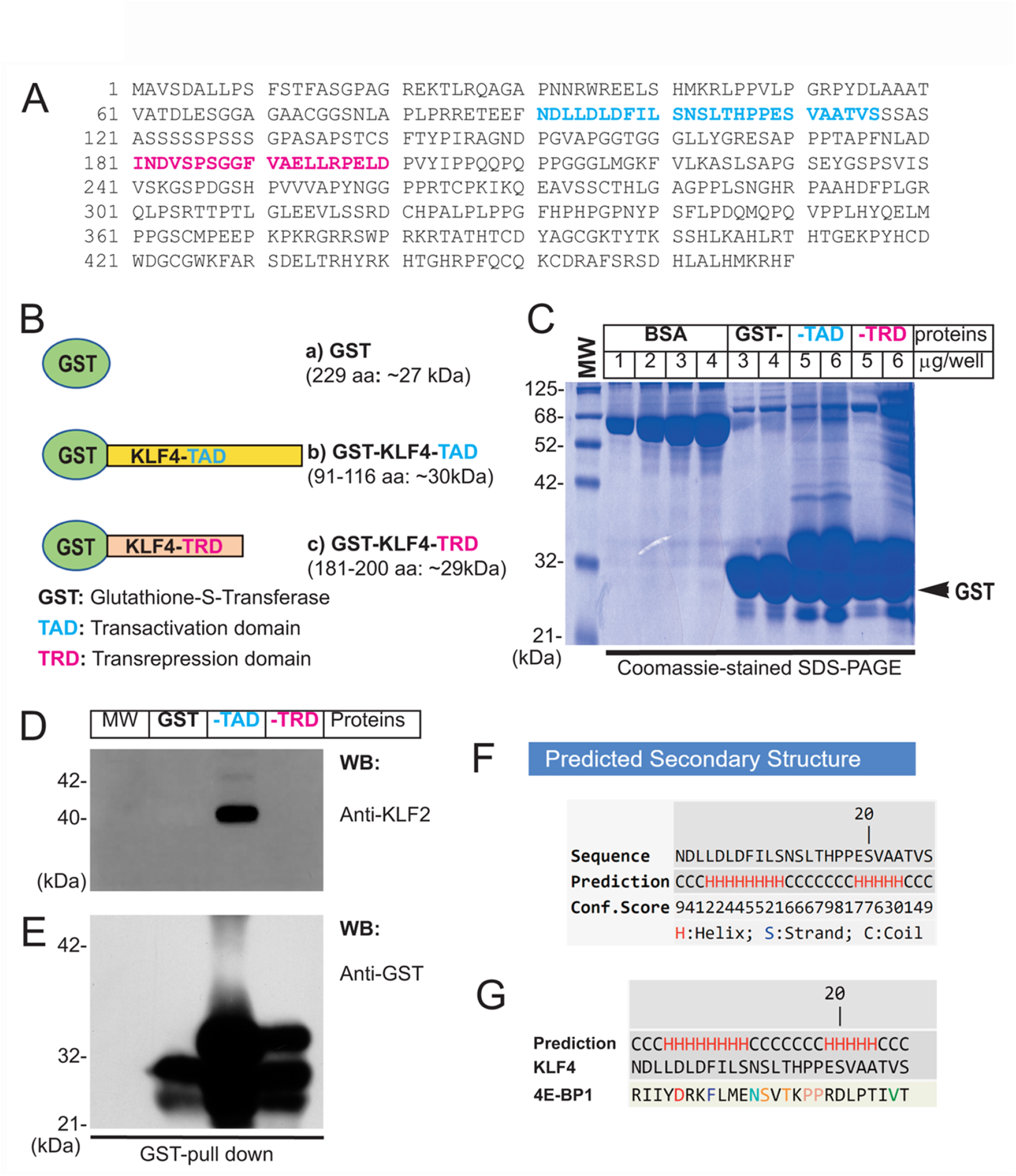
Direct interaction of KLF4 with KLF2. **A.** Amino acid sequence of human KLF4. The amino acid (TAD) sequence 91-116 is shown in blue color, while magenta represent TRD region used for preparation of GST-fusion proteins. **B**. Schematics of GST-fusion protein constructs: a) GST-alone (∼27 kDa), b) GST-KLF4-TAD (∼30 kDa), and c) GST-KLF4-TRD (∼29 kDa). C. GST-fusion proteins were expressed in *E coli* (BL2) and purified. **C**. The integrity of GST-fusion proteins was determined by SDS-PAGE. **D**. hLMECs nuclear extracts were precleared and pre-absorbed with ∼5.0μg fusion protein-Sepharose beads; thereafter, 10mg/nuclear protein extracts were incubated with indicated sepharose-4B bound-GST-fusion proteins (3.0μg/GST-fusion protein construct) for 30 minutes at 4°C, washed 5 times with nuclear extraction buffer, and resolved by SDS-PAGE. The NC membrane was probed with anti-KLF2 antibody. **E**. The membrane was stripped and reprobed with anti-GST antibody to examine equal loading of GST-fusion proteins across the lanes. Experiments were performed at least 3 times. **F&G**. KLF4-TAD amino acid sequence, showing helix and coiled-coil secondary structures performed using I-TASSER online computational analytical tools.

### 4. Endothelial *Klf4*-knockdown promotes upregulation of Klf2

Close examination of ∼1.0 kbp upstream of the transcription start site (TSS) in the human *KLF2*-promoter/enhancer showed the presence of six putative KLF4/KLF2 binding sites, 5’-CACCC-3’ in forward and 5’-GGGTG-3’ in reverse orientation (Fig. 4A). The DNA sequence of the human *KLF2*-promoter/enhancer segment is shown in Fig. 4B. We thus designed oligonucleotide PCR-primers to amplify the flanking six putative KLF4/KLF2 binding sites on the *KLF2*-promoter DNA sequence (Fig. 4C). Then, hLMVECs at 70-80% confluence were treated with *control-shRNA* and *KLF4-shRNA* to induce knockdown. Thereafter, sheared chromatins were immunoprecipitated with indicated antibodies and eluants were PCR-amplified (Fig. 4D). The primers amplified a 487 bp PCR-product suggesting thee was binding of KLF4 to the *KLF2*-promoter, while IgG and KLF2 did not. However, in *KLF4-shRNA* treated hLMVECs, KLF2 bound to its own promoter (middle lane), while IgG and KLF4 did not (Fig. 4D). The efficiency of knockdown was analyzed by WB with indicated antibodies (Fig. 4E). This data suggests that at homeostatic levels, KLF4 binds to the *KLF2* promoter, while loss or decreased expression of KLF4 allows KLF2 to auto-regulate its own expression by binding to its own promoter. To confirm this hypothesis, we performed an electrophoretic mobility shift assay (EMSA). Biotin-labeled oligonucleotide probes (P1, P2, and P3) designed based on the *KLF2*-promoter harboring 5’-CACCC-3’ or 5’-GGGTG-3’ DNA sequences were used for the EMSA. After incubation with nuclear extracts, biotin-labeled oligonucleotide probe interaction was observed between KLF4 with P1, with the super-shifts indicated by colored arrows (Fig. 4G). In a converse experiment, EMSA was performed using KLF4-depleted nuclear extracts (Fig. 4H); the result showed that in absence of KLF4, KLF2 binds to the same oligonucleotide probe (P1), indicating KLF4 and KLF2 are able to occupy the same exact DNA sequence.

**Figure 4.**
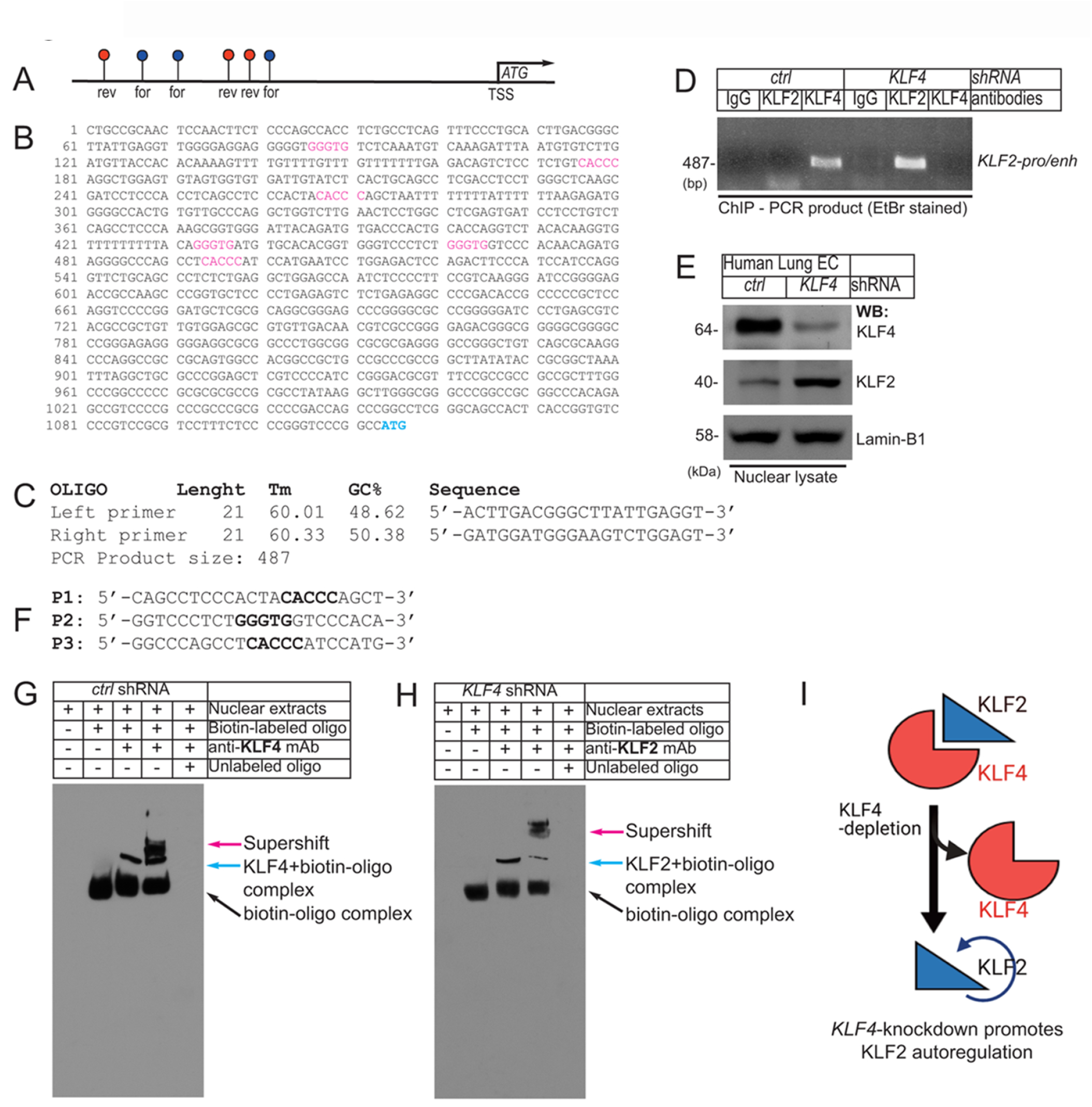
EC Klf4-depletion promotes Klf2 autoregulation. **A**. Schematics of ∼1.0 kbp upstream of the human *KLF2*-promoter/enhancer shows presence of 6 putative KLF4/KLF2 binding sites (blue, in forward-; red, in reverse-orientation). **B**. DNA sequence of the human *KLF2*-promoter/enhancer segment. The putative KLF4/KLF2 binding sites are as shown (For-CACCC and Rev-GGGTG). ATG indicates transcription start site (TSS). **C**. Oligonucleotide PCR-primer sequences used for amplifying KLF4/KLF2 binding sites. The PCR primers produce ∼487 bp PCR-product. **D**. Chromatins were prepared from hLMVECs that were subjected to *control-shRNA* and *KLF4-shRNA* knockdown. Chromatins were sheared, clarified, and pre-absorbed with Sepharose-protein-A bound IgG (2.0μg/ChIP) for one hour at 4°C. Thereafter, immunoprecipitated with indicated antibodies (2.0μg/ChIP) and eluants were PCR-amplified by primers. The PCR-product ∼487 bp indicates binding of KLF4 to cognate promoter/enhancer region harboring - CACCC- and -GGGTG- in the KLF2 promoter. In hLMVECs, subjected to *KLF4-shRNA,* KLF2 bound to its own promoter. **E**. Efficiency of shRNA mediated knockdown are as shown. **F**. Biotin-labeled oligonucleotides (P1, P2, and P3) designed after the *KLF2*- promoter DNA sequence used for EMSA. **G&H**. Biotin-labeled oligonucleotide (P1) harboring the sequence (CACCC) of the putative KLF2/KLF4 binding sites designed after the *KLF2*-promoter were incubated with nuclear extracts. The supershift was carried-out by preincubation (30 minutes) with excess nuclear extracts, prepared from **G**) control-shRNA or **H**) *KLF4*-shRNA knockdown ECs with indicated antibodies. Experiments were repeated 3 times. **I**. Summary model showing the loss of KLF4 promoted binding of KLF2 to the *KLF2-*promoter/enhancer DNA sequence.

### 5. *Klf4*-deletion in quiescent ECs propagates an EndMT phenotype

To address the *in vivo* role of KLF4 in EC quiescence, we bred and crossbred several different strains of mice to produce: (1) *Klf4^fl/fl^::Cdh5^CreERT2^* and (2) *Rosa^mT/mG^::Klf4^fl/fl^::Cdh5^CreERT2^* mice lines in C57BL background (Fig. S4A-D). The *Cdh5*-promoter driven CreERT2-mediated genetic deletion following TAM administration allowed EC-lineage tracing (Fig. 5A&B). Next, CD45- and CD31+ ECs (see methods) from the control and TAM-administered mice were used for biochemical analyses. Accordingly, WB analyses of cell extracts prepared from CD45- and CD31+ lung ECs isolated from these mice confirmed Klf4-silencing (henceforth called *Klf4*^ECKO^). In *Klf4^ECKO^* mice, we observed increased Klf2 expression (Fig. 5C), increased TGF-β1, increased type-I collagen protein levels, and phosphorylation of Smad3 (Ser465/467), but decreased VE-cadherin (Fig. 5C). Together, these data strongly suggest that *Klf4*-silencing is associated with increased Klf2-expression and EndMT.

**Figure 5.**
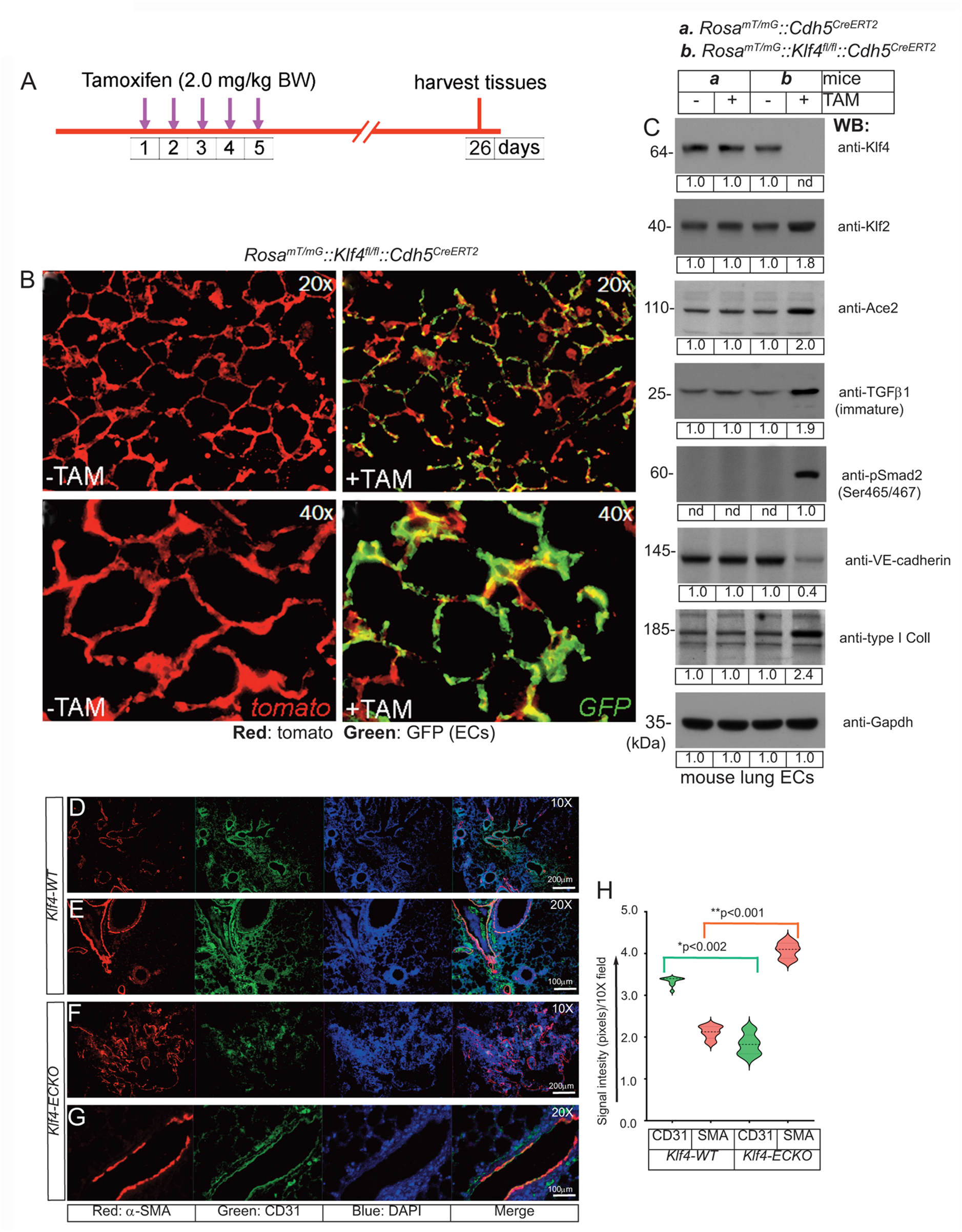
EC-*Klf4* deletion induces EndMT in the lung tissues. **A**. The timeline and TAM treatment scheme; **B**. Mice are injected with 2.0mg/kg body weight TAM for five consecutive days to induce deletion of EC-*Klf4* alleles and allow EC-GFP expression. In absence of TAM, all cells in reporter mouse express tomato (red) fluorescence. Lung tissue samples are harvested on day 26. Magnifications are as shown. **C**. EC extracts were prepared from indicated mouse lung tissues and analyzed by WB with indicated antibodies. Molecular weights are given in kiloDalton (kDa). The numbers below each WB panel indicate signal quantification. Experiments were performed at least 3 times. **D-G**. Immunohistological analysis of lung tissues prepared from *Klf4-WT* (-TAM) and *Klf4- ECKO* (+TAM) mice (*Klf4-WT*, n = 10; *Klf4-ECKO*, n = 10) with anti-CD31 and anti-α- SMA. Note that increased anti-α-SMA antibody immunoreactivities (red) exclusively co-aligned with CD31+ vascular structures. Nuclei were counterstained with DAPI (blue). Magnification of images are as shown. Experiments are done in both male and female mice (5 + 5) and the EndMT pathology were identical. Magnifications and scale bars are as shown. **H**. Quantification of red (SMA) and green (CD31) signal intensities using NIH Image-J software in arbitrary units; *P < 0.002; **P < 0.001 compared with Klf4-WT; unpaired 2-tailed Student’s t test. At least eight 10X microscopic fields/per slide were used for quantification and 5 slides were used for each group. Staining experiments were repeated at least 3 times and examined under Olympus Epifluorescence microscope in room temperature. Images were saved as raw data, converted into TIFF format and combined using QuarkXpress 2020 software, finally saved as TIFF documents.

To further address the underlying relationship between the loss of quiescent EC-Klf4 and EndMT of lung ECs, we monitored the effect of *Klf4*-deletion in *Klf4*^ECKO^ mice (Fig. 5D-H). Four weeks after the last TAM injection, microscopic examination of histological tissue sections showed a profound increase in anti-αSMA antibody immunoreactivity of CD31+ vascular structures in *Klf4-ECKO* lung tissues (Fig. 5F&G). Quantification showed a >50% decrease in CD31+ signal intensity (Fig. 5H), while anti-αSMA immunoreactivity increased significantly (Fig. 5H). These data indicate that in *Klf4*-ECKO lung tissues, ECs acquire the *bona fide* mesenchymal marker α-SMA, consistent with the WB data shown in Fig. 5C indicating EC-*Klf4*-silencing is strongly associated with EndMT in *Klf4*-ECKO.

### 6. *Klf4*-deletion in ECs leads to pulmonary fibrosis

Recent studies suggest that EndMT-derived myofibroblasts are one of the main drivers of pulmonary fibrosis. To explore the potential connections among the loss of EC-Klf4, EndMT, and lung pathology, tissue sections were subjected to Masson’s trichrome stain. The timeline and strategy are shown at the top (Fig. 6). For this experiment, 10 lung lobes were collected from five (5) *Klf4*-WT control and five (5) *Klf4*-ECKO mice. Microscopic examinations were carried-out at 10x, 20x and 40x magnifications (Fig. 6A-L) and NIH-Image-J software was used for quantification. *Klf4*-WT lungs showed normal alveolar architecture, while in *Klf4*-ECKO mice the alveolar architecture appeared distorted with an increase in fibroblast density and trichrome+ (blue color) collagen deposition (Fig. 6D-F&J-L) which was quantified as shown Fig. 6M. Collagen deposition characterized by trichrome staining was consistent with the WB analysis shown in Fig. 5C. Therefore, these data indicate that EC-*Klf4* deletion is responsible for lung fibrosis observed in *Klf4*-*ECKO* mice.

**Figure 6.**
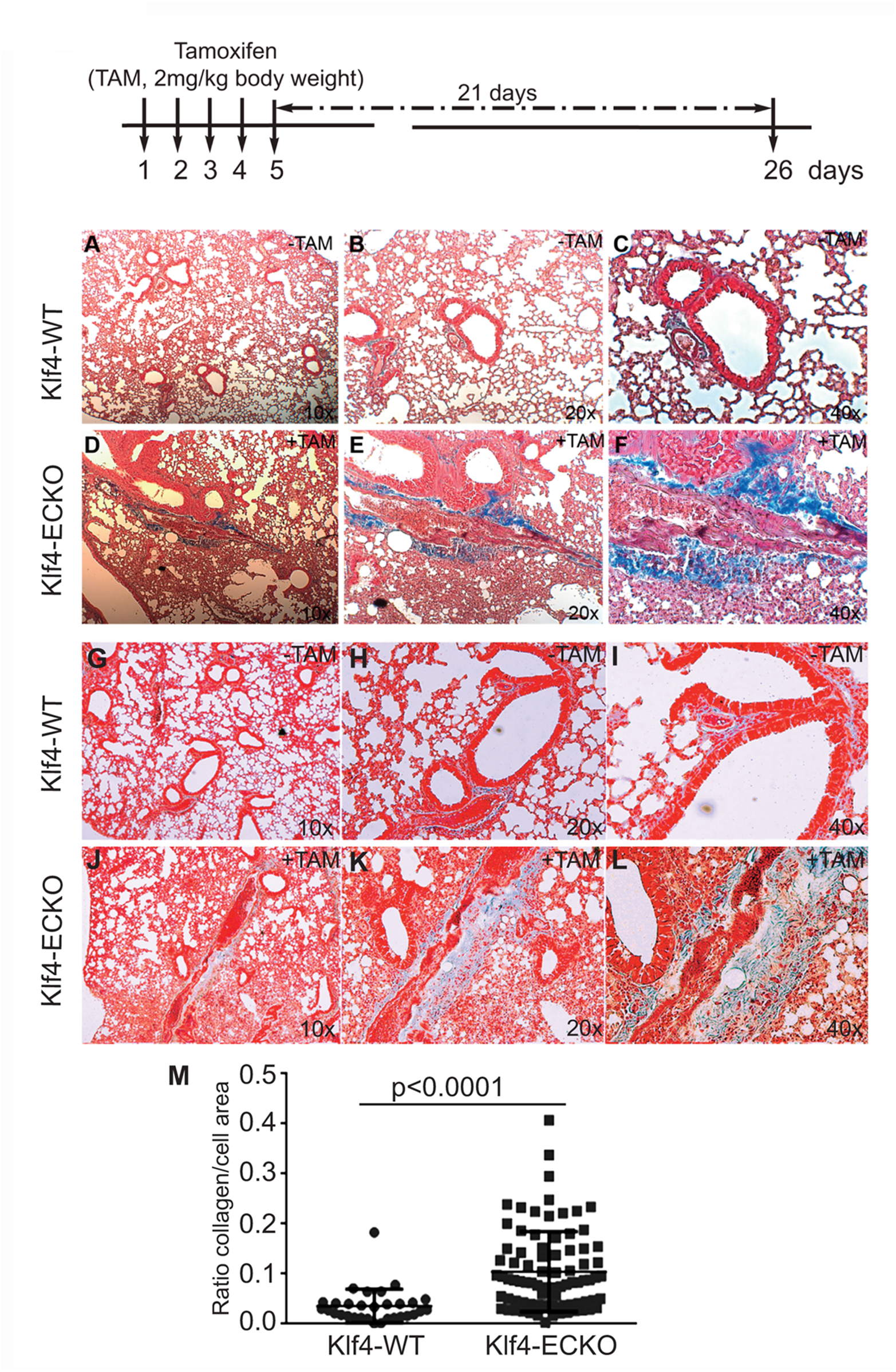
EC-*Klf4* deleted mice harbor appreciable level of fibrosis with alveolar enlargement. Top, strategy and timeline of TAM injection experiments. At the end of experiments (day 26), 8 lungs from control and 8 from *Klf4*-ECKO mice were excised and prepared for histology by fixing and staining with Masson’s Trichrome. All microvessels in pulmonary cross-sections were imaged on a Zeiss Axiovert microscope at 20X magnification. For blinded measurements, NIH-Image J software was used. Representative images of hematoxylin and eosin (H&E) stained cross-sections of mouse lung showing representative lung architecture. **A-L**. Normal architecture of alveolar and capillaries in *Klf4*-WT and Klf4-ECKO mice at indicated magnifications (10x, 20x, and 40x). **M**. Quantification of collagen (blue color) staining Kruskal–Wallis test followed by Dunn’s multiple comparisons to Klf4-WT mice. Data are mean ± S.D. n = 8; *p<0.0001 vs. Klf4-WT (corn oil).

### 7. Mice harboring EC-*Klf4* deletion show hallmarks of EndMT

To further explore the extent of EndMT, primary CD31+/CD45- mLMVECs were isolated from 14 weeks old male and female Klf4-WT and Klf4-ECKO mice. The strategy and timeline of the experiment is shown in Fig. 7A. Please see methods for CD31+/CD45- isolation procedures. Total protein extracts were subjected to WB analyses with anti-VEGFR2/Flk1, anti-VE-cadherin, anti-VCAM-1, anti-Klf4, anti-Klf2, anti-p-Smad3 (S423/425), and anti-α-SMA antibodies (Fig. 7B). The expression of VEGFR2/Flk1 did not change in these ECs. Compared to control, there was decreased expression of VE-cadherin in Klf4-ECKO ECs, while EC-Klf2 and VCAM-1 increased. Importantly, the expression of EC-Klf2 and VCAM-1 increased in the Klf4-ECKO cohort. Additionally, we observed increased phosphorylation of Smad3 (S423/425) and acquisition of α-SMA expression in ECs of Klf4-ECKO mice. GAPDH was used to demonstrate equal loading of protein in each lane (Fig. 7B). Endothelial cell-specific characteristics of CD31+/CD45- cells isolated from Klf4-WT and Klf4-ECKO was assessed by anti-VE-cadherin staining (Fig. 7C-F). Compared to control, ECs isolated from Klf4-ECKO showed decreased VE-cadherin staining by epifluorescence microscopy Fig. 7E&F. These data indicate loss of EC-Klf4 is strongly associated with EndMT.

**Figure 7.**
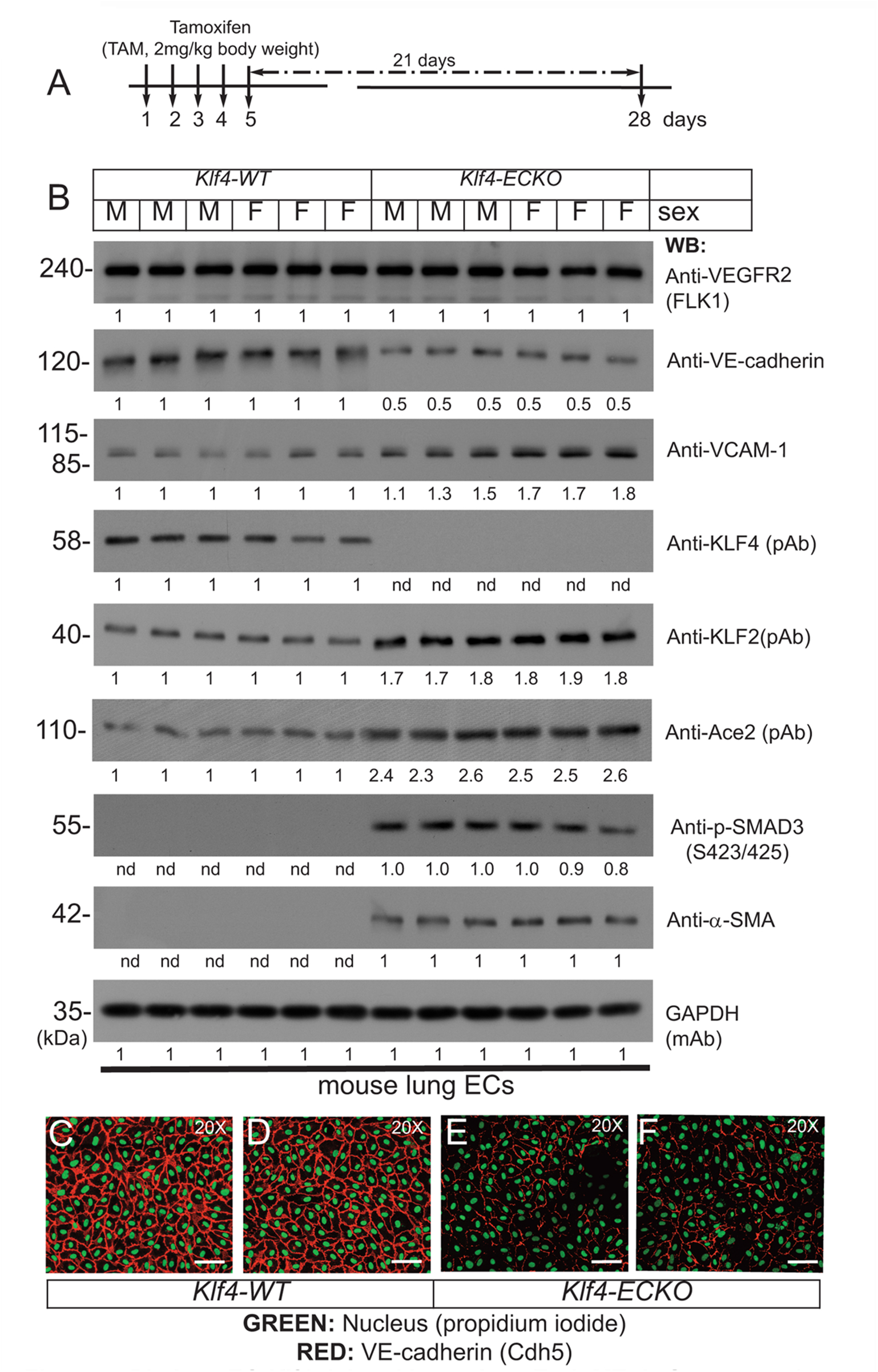
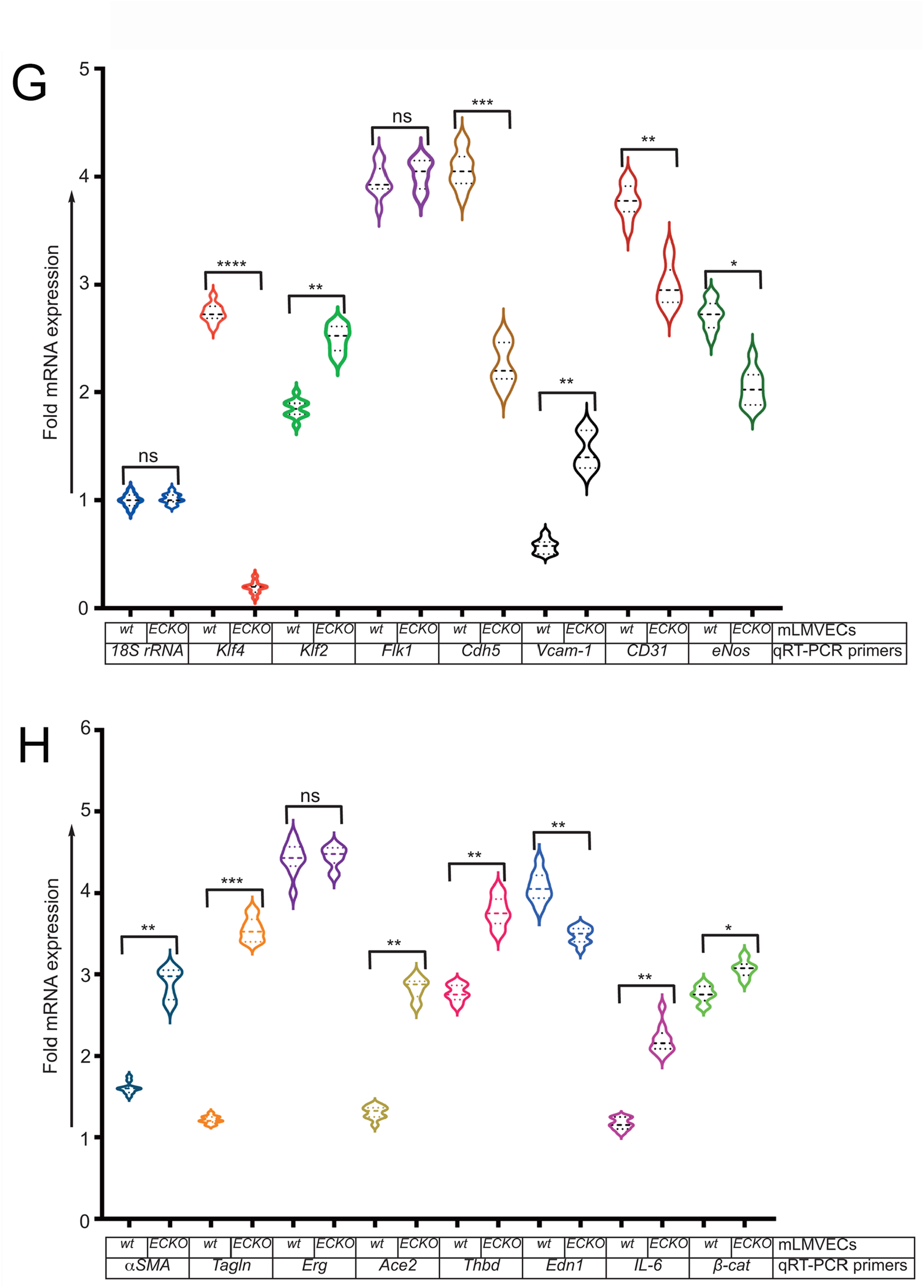
EC-*Klf4* deleted lung show EndMT. **A**. Strategy and timeline of experiments. **B.** CD31^+^/CD45^−^ lung microvascular EC protein extracts prepared from 12-14 weeks old male (M) and female (F) *Klf4*-WT (corn oil) mice and *Klf4*-ECKO (+TAM) mice were analyzed by WB with indicated antibodies. VEGFR2 is expressed in all mice with no apparent change in the level of expression. The expression of VE-cadherin in ECs was decreased, while the expression of EC-Klf2, Ace2 and VCAM-1 increased, in ECs of *Klf4*-ECKO mice. We also observed increased phosphorylation of SMAD-3 (S423/425) and acquisition of α-SMA in all ECs that lacked Klf4. There was no change in Gapdh levels. Experiments were repeated at least 3 times. The numbers (quantification) below the WB panels show relative signal intensities. **C-F.** Representative images of EC characteristics of mouse lung ECs were determined by anti-mouse VE-cadherin (Texas red) staining; nuclear Propidium iodide (green). Magnification, 20×; scale bar, 250 μm. **G&H**. Total mRNAs prepared mLMVECs isolated from Klf4-wt and Klf4-ECKO mice were analyzed by qRT-PCR as described in methods. 18S rRNA was used as control. Data represent mean ± S.E.M.; ns, not significant; *p<0.5; **p<0.01; ***p<005; ***p<0.001 versus Klf4-wt as shown. Experiments were repeated 3 times with triplicates.

Next, mRNA prepared from *Klf*-WT and *Klf4*-ECKO mLMVECs was analyzed by qRT-PCR assay with mouse gene specific oligonucleotide primers (Table 2). The expression of control *18S rRNA* did not change, while *Klf4* expression decreased significantly in mLMVECs obtained from *Klf4*-ECKO mice (Fig. 7G). The expression of *Klf2* increased significantly (p<0.001) in *Klf4*-ECKO group, but Flk1/Vegfr2 did not (Fig. 7G). In addition, the expression of several EC-defining genes were decreased in *Klf4*-ECKO mLMVECs, such as *Cdh5* (*VE-cadherin;* ∼ 1.5 fold), as well as *CD31*, *Edn1*, and *eNos* (Fig. 7G&H). Importantly, the EndMT markers α*SMA* and *Tagln* (*sm22*) and inflammation markers *Vcam-1* and *IL-6* increased significantly, whereas *Erg* did not (Fig. 7G&H). We also observed significantly increased expression of *Ace2* and *Thbd* in the *Klf4-*ECKO group, as well as a moderate increase of β-catenin expression signifying ongoing EndMT (Fig. 7H). These data show that loss of EC-Klf4 is strongly associated with EC dysfunction and EndMT.

### 8. EC-Klf4 deletion is associated with alveolar enlargement and increased immune cell adhesion to the endothelium

As we observed extensive EndMT in Klf4-ECKO mouse lung tissues, we next explored whether Klf4 gene deletion was associated with disruption of lung alveolar structures. The timeline and strategy of this experiment is shown in Fig. 8A. Cohorts of Klf4-WT and Klf4-ECKO mouse lung tissues were stained with H&E (Fig. 8B-F). Detailed morphometric analyses showed no differences in alveolar structures among control mice (Fig. 8B-D). In contrast, Klf4-ECKO mice showed widespread alveolar collapse (Fig. 8E&F). Quantification revealed a significant decrease in the number of alveoli (Fig. 8F&G). We also observed fluid-filled alveoli, sequestration of neutrophils, and degradation of basement membrane in Klf4-ECKO cohorts indicating the presence of vascular inflammation and injury associated with an alveolar pathology (Fig. 8E-G). These data indicate that the loss of EC-Klf4 results in altered alveolar structural integrity.

**Figure 8.**
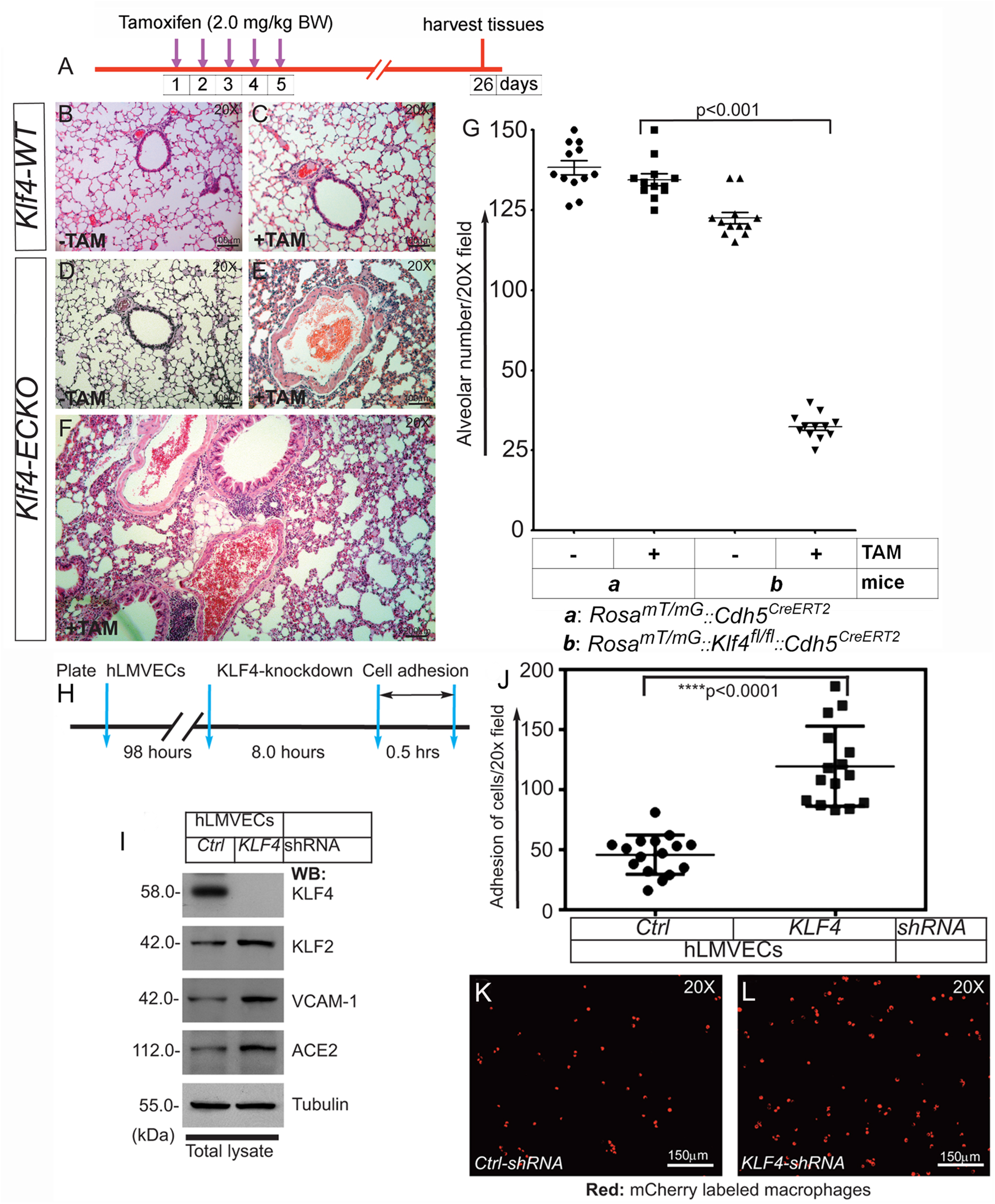
EC-*Klf4* deleted mice harbor decreased number of alveoli and enlarged alveolar structures. **A. Strategy and timeline of experiments. B&C.** H&E staining show no change in alveolar structures in control and TAM treated cohorts of Klf4-WT mice. **D&E.** While there are no changes detectable in Klf4-ECKO (-TAM) cohort, however, there are significant changes accompanied by cellular infiltration and changes in alveolar architecture in a cohort of Klf4-ECKO mice (+TAM). **F**. A magnified image of lung pathology in a Klf4-ECKO mice. **G**. Quantification of alveolar structures. P < 0.001 compared with Klf4-WT (+TAM) group. n = 12 mice (6 male and 6 female mice) unpaired 2-tailed Student’s t test. Experiments were repeated 2 times. **Macrophage adhesion onto confluent human lung microvessel ECs (hLMVECs) monolayer. H.** Knockdown, strategy, and timeline of experiment. hLMVECs were treated with retroviral particles encoding *ctrl-shRNA* or *KLF4-shRNA* for 8hrs. **I**. Efficiency of knockdown was determined by WB with indicated antibodies. Molecular weights are shown in kDa. **J**. Next, fluorescent mCherry labeled RAW 264.7 macrophages were allowed to adhere onto hLMVECs for 0.5 hrs. Next, hLMVECs were washed to remove non-adherent cells and macrophage adhesion events were quantified by counting the number of fluorescent mCherry/red macrophages per 20X microscopic field. Quantification macrophage adhesion onto hMLECs in response to KLF4-depletion. n = 16-18; ****P<0.0001 vs control shRNA. Statistics were carried out using 1-way ANOVA with Newman-Keuls post-hoc test for multiple comparison correction. **K&L**. Representative confocal images of mCherry (red)-labeled macrophages onto hLMVECs. Experiments were repeated at least three times.

Pro-inflammatory cytokines including TGF-β1, TNF-α, and IL-6 increase expression of cell surface proteins including E-selectin, ICAM, and VCAM-1. These cell surface adhesion molecules recruit and promote firm adhesion of immune cells to injured or activated ECs. Given that Klf4-ECKO was associated with increased expression of VCAM-1 and ACE2, recruitment of inflammatory cells, and extensive EndMT in mouse lung tissues, we next asked the question if EC-KLF4 depletion makes the ECs more adherent to immune cells. To address this question, we performed a cell adhesion assay on KLF4 knockdown ECs (Fig. 8H-L). The efficiency of knockdown was determined by immunoblotting (Fig. 8I). In the cell adhesion assay, we observed increased attachment of macrophages to monolayer-ECs that received KLF4-shRNA compared to control shRNA treated ECs (Fig. 8J-L). This finding indicates that loss of EC-KLF4 increases expression of cell adhesion molecules which bind to counterreceptors such as VLA-4, also known as integrin α_4_β_1_, that are abundantly expressed on the surface of immune cells such as macrophages.

## Discussion

Quiescent ECs crucially maintain lung alveolar homeostasis in adults (7,8). These cells continuously sense humoral, paracrine, and angiocrine factors and mechanical forces, but are also able to preserve the integrity of the airway microenvironment (1–4). Our understanding of the molecular mechanisms central to EC quiescence should provide clues to repair mechanisms for EC-dysfunction, normalization of tumor blood vessels, and potential therapies to reverse the course of EndMT. Accordingly, we showed that quiescent ECs express KLF4 and KLF2 proteins which were found bound to each other as a heterodimeric protein complex. However, in the event of a decrease in KLF4 level, KLF2 overcompensated for the loss of KLF4 by upregulating its own expression which led to the disruption of KLF4-mediated quiescent EC phenotype and propagated the emergence of EndMT with alveolar enlargement and fibrosis.

In healthy lung tissues, KLF2 and KLF4 can be detected, however, the expression patterns of KLF4 and KLF2 changed dramatically in diseased lungs. Specifically, the expression of KLF4 decreased, while KLF2 increased significantly in diseased human lung tissues. Quantification showed that KLF4 expression was barely detectable in lungs from patients with emphysema and COPD, while the expression level of KLF2 clearly increased. In addition, we observed increased ACE2 and decreased VE-cadherin in these tissue samples. As TGF-β1 is frequently associated with fibrotic disease, we also addressed TGF-β1 levels in these tissue samples and observed an appreciable increase in the expression of TGF-β1. Since we had access to a very limited number of lung tissue samples, the mechanistic relationship among these proteins and causality of these human diseases is not possible from the data presented. However, increased expression of TGF-β1 strongly corelates with the occurrence fibrosis and thus these data provided the impetus to develop an inducible mouse model test the hypothesis that the oscillations in KLF4 and KLF2 expression might play a central role in the induction of EC dysfunction, EndMT, and fibrosis.

Biochemical analyses of quiescent EC nuclear extracts showed the presence of mostly monomeric, but also di- and tetrameric forms of KLF4 and KLF2 polypeptide species. These oligomeric complexes were observed in the absence of reducing agent (β-mercaptoethanol), whereas the addition of 5% β-mercaptoethanol dissociated the oligomers, thereby producing monomeric forms of KLF4 and KLF2. These results suggest that quiscent ECs harbor mostly monomeric-KLF4 and -KLF2 proteins, but to a lesser extent, dimeric and tetrameric forms can also be found in these cells. One could surmise that there may be more than one pool of KLF4 or KLF2 proteins, e.g., KLF4-monomers KLF4-KLF4 homodimer, and KLF4-KLF4 homotetramer. In addition, reciprocal immunoprecipitation experiments indicated existence of KLF4-KLF2 heterodimeric complexes of proteins that included p300. A more complex interpretation could that the quiescent ECs are heterogeneous and the existence of mono-, di- or tetrameric forms of KLF4 and KLF2 might indicate the degree of cellular differentiation or cellular maturation states. In other words, the KLF4-KLF2 complexes might be present in only a subset of quiescent ECs. However, we cannot rule out the possibility that presence of various complexes of KLF4 and KLF2 might also be indicative of different phases of cell cycle. Nevertheless, KLF4 and KLF2 are expressed in quiescent ECs and these two proteins are known to associate directly with p300, a nuclear protein that acts as a histone acetyltransferase (HAT) to transcriptionally regulate targets through chromatin remodeling (34–36). That is, p300 acetylates KLF4 to regulate gene transcription via the modulation of histone acetylation. KLF2, KLF4, and p300 likely form a multimolecular complex in quiescent ECs, consistent with the known ability of transcription factors such as c-Myc to dimerize with Max, Fos with Jun, and β-catenin with TCF4/LEF1. STAT1 homodimerizes with STAT1, and STAT1 heterodimerizes with STAT5. These physical associations are often meaningfully regulated by upstream signaling elements that induce the expression of downstream target genes which could mediate cellular responses such as cell differentiation, proliferation, migration, apoptosis, and survival. However, the mutation, epigenetic modification, or overexpression of transcription factors is associated with pathology. For example, c-Myc and β−catenin mutations are frequently found in many tumors. Importantly, we characterized the amino acid sequence of KLF4 that mediated direct binding to KLF2 to form a heterodimeric proteins complex. Further analyses revealed these amino acid residues of KLF4 that can form α-helical and coiled-coil secondary structures, and which could form a 3D structure analogous to eukaryotic translation initiation factor 4E (eIF4E)-binding protein 1 (4E-BP1). 4E-BP1 belongs to a family of translation repressor proteins and is a substrate of mTOR signaling cascade, however, it’s function in quiescent ECs remains incompletely understood.

Expression analyses performed by several laboratories has demonstrated that KLF4 and KLF2 share overlapping gene targets—about 40% of gene targets are coordinately regulated by both KLF4 and KLF2. The ability of KLF4 and KLF2 to induce expression of a subset of EC gene promoters is not surprising since KLF4 or KLF2 are both known to bind to promoter and enhancer segments harboring 5’-CACCC-3’ or 5’- GGGTG-3’ DNA sequences. However, they can also induce the expression of specific genes in a non-overlapping manner; therefore, most studies have addressed the ability of KLF4 or KLF2 to act alone. While KLF2 and KLF4 can induce the expression of genes independently, no studies to date have addressed the ability of KLF4 to bind to KLF2 or regulate its expression. Thus, we determined which segment of KLF4 interacts with KLF2. To address this question, we conducted mutational analyses combined with transfection experiments. Nuclear extracts prepared from hLMVECs expressing equivalent levels of flag-epitope tag constructs showed reciprocal co-IP of KLF4 bound to KLF2 through the 97–117 amino acid residues present within the KLF4-TAD segment. Interaction of KLF4 with KLF2 was also shown by GST pull-down which revealed that KLF4-TAD interacts with KLF2. In other words, KLF4 heterodimerizes with KLF2.

To clarify the exact relationship between KLF4 and KLF2, we examined approximately -1.0 kbp upstream of the TSS of the human KLF2-promoter/enhancer. Interestingly, we detected six putative KLF4/KLF2 binding sites in the KLF2promoter/enhancer segment. This finding could potentially explain why a decrease in KLF4 level might induce the upregulation of KLF2. Our data show that, in control-quiescent ECs, KLF4 bound to the KLF2 promoter, and that a decrease in KLF4 expression was associated with KLF2 autoregulation of its own expression by binding to its own promoter. The ability of KLF4 and KLF2 to bind to the same target sequence was further confirmed by an EMSA experiment. In a converse experiment using KLF4-depleted nuclear extracts, we observed KLF2 binding to the same target sequence indicating KLF4 and KLF2 occupy the same exact DNA sequence. This suggests that KLF4 and KLF2 are intrinsically connected and may regulate lung EC gene expression collaboratively and co-operatively.

Next, we addressed the role of KLF4 in ECs *in vivo* by generating *Klf4^fl/fl^::Cdh5^CreERT2^* and *Rosa^mT/mG^::Klf4^fl/fl^::Cdh5^CreERT2^* mice. We used both female and male mice that were 12–14 weeks old for experiments and applied a previously established protocol of low-dose TAM (2.0mg/kg body weight) injected intraperitoneally (i.p.) for five consecutive days, and on day 28, studies were conducted. We used EC-lineage methodology to label ECs and sorted GFP+ ECs for our experiments. Microscopy and WB analyses confirmed the EC-specific loss of Klf4 (called *Klf4*-ECKO). Importantly, we detected increased EC-Klf2 expression in *Klf4*-ECKO lung ECs. In *Klf4*-ECKO cells, we also found decreased VE-cadherin expression, but increased Ace, TGF-β1, type-I collagen, and phospho-Smad3 (Ser465/467). A decrease in VE-cadherin signifies the loss of adherens junction mediated cell−cell adhesion, a hallmark of EndMT, while the increase in TGF-β1 and collagen expression are indicative of a pro-fibrotic pathway. Taken together, these data revealed that loss of EC-Klf4 is associated with increased Klf2 expression and EndMT. Furthermore, we detected a profound increase in the anti-αSMA antibody immunoreactivity in the CD31+ vascular structures in *Klf4*-ECKO lungs. Microscopy clarified the co-alignment of CD31+ ECs with anti-α-SMA positivity. To explore further the mechanistic link between loss of EC-Klf4, EndMT, and lung pathology, tissue sections were analyzed by Masson’s trichrome staining which revealed damaged and distorted alveolar architecture in *Klf4*-ECKO mice. There were also indications of immune cell infiltration, enlarged alveoli, and deposition of collagens. Together, these data strongly indicated that the loss of EC-Klf4 is associated with lung disease.

Furthermore, biochemical analyses of proteins prepared from *Klf4*-ECKO mice showed decreased VE-cadherin, but increased Klf2, VCAM-1, Ace2, phospho-Smad3 (S423/425), as well as increased αSMA levels. Image analyses confirmed the decrease in VE-cadherin staining at cell-cell junctions of endothelial cells lining lung microvessels. In addition, qRT-PCR experiments confirmed the increase in expression of *Klf2*, *Vcam-1*, *IL-6*, *α-SMA*, *Tagln*, *Thbd*, and *β-catenin*, while *Klf4*, *Cdh5*, *CD31*, and *eNos* mRNA decreased, signifying EC-dysfunction and ongoing EndMT. Thus, these data reveal the loss of EC-Klf4 is strongly associated with EndMT. In addition, we detected neutrophils and degradation of basement membrane in *Klf4-*ECKO cohorts indicating the presence of EC dysfunction and alveolar damage. Therefore, these data illustrate that the loss of EC-Klf4 is incompatible with the normal alveolar structural integrity.

It is increasingly clear that immune cells adhere to the endothelium in response to inflammation, injury, and regenerative processes. As Klf4-deficient ECs showed increased expression of Klf2, VCAM-1, ACE2, and IL-6, we addressed whether the endothelium loses its anti-inflammatory and anti-adhesive properties. As expected, KLF4 knockdown increased adhesion of macrophages onto monolayer ECs. Specifically, we observed increased attachment of macrophages to EC monolayers that received KLF4-shRNA. This indicated that the loss of EC-KLF4 promotes the expression of cell adhesion molecules such as VCAM-1, which then can bind to a counter-receptor such as VLA-4 (aka α_4_β_1_ integrin) expressed on monocytes/macrophages. Together, these findings are consistent with the known function of KLF4 to act as an anti-inflammatory and atheroprotective transcription factor (32,36).

## Conclusion

KLF4 and KLF2 are highly expressed in quiescent ECs. However, EC-specific inducible *Klf4*-deletion induced irreversible EC activation and EndMT. Loss of KLF4 was accompanied by an increase in KLF2 expression, increased VCAM-1, ACE2, and IL-6 expression, and recruitment of circulating immune cells. An emerging hypothesis that stems from these observations is that the loss of quiescent EC-KLF4 is likely pathogenic, therefore normalization lung microvessel KLF4 levels in patients with chronic lung disease might reverse the course of EndMT and restore the EC quiescence (please see graphical abstract).

## Methods and Materials

### Antibodies, Cell-Molecular Biology and Biochemical Reagents

Goat polyclonal anti-m/rCD31 (AF3628) was purchased from R&D Systems (Minneapolis, MN). Mouse monoclonal anti-KLF4 (4E5C3) (200-301-CE9) and anti-Lamin-B1 were purchased from Rockland Immunochemicals, Inc (Limerick, PA). Mouse monoclonal anti-α-smooth muscle actin (α-SMA) (A2547), and RNAzol RT (R4533) were purchased from Sigma-Aldrich Corporation (St. Louis, MO). Rabbit polyclonal anti-TAGLN/Transgelin (ab14106), sheep polyclonal anti-von Willebrand Factor (ab11713), rabbit polyclonal anti-Collagen I (ab34710), donkey polyclonal anti-sheep Immunoglobulin G (IgG) Alexa Fluor 488 (ab150177), and donkey polyclonal anti-rabbit Immunoglobulin G (IgG) Alexa Fluor 647 (ab150063) were purchased from Abcam (Cambridge, MA). Rabbit polyclonal anti-KLF4 (H-180) (sc-20691), mouse monoclonal anti-Klf4 (F-8) (sc-166238), rabbit polyclonal anti-KLF2 (H-60) (sc-28675), mouse monoclonal anti-GAPDH (4G5) (sc-51906), mouse monoclonal anti-human p300 (NM11) (sc32244), mouse monoclonal anti-TGF-β1 (sc130348), goat polyclonal anti-human β-tubulin (E-14) (sc86255), goat polyclonal anti-human VE-Cadherin (CDH5) (C-19) (sc6458), mouse monoclonal anti-Endoglin (A-8) (sc376381), mouse monoclonal anti-Endoglin (P3D1) (sc18838) were purchased from Santa Cruz Biotechnology (Santa Cruz, CA). Rat anti-mouse CD31 (550274) was purchased from BD Biosciences (San Jose, CA). Rabbit anti-mouse von Willebrand Factor (vWF) (AB7356) were purchased from EMD Millipore (Billerica, MA). Rabbit monoclonal anti-P-Smad3 (S465/467) (3108S), rabbit monoclonal anti-alpha-Smooth muscle actin (D4K9N) XP (19245S), and rabbit monoclonal anti-human VE-Cadherin (D87F2) XP (R) (cat#2500S) were purchased from Cell Signaling Technology (Danvers, MA). Mouse monoclonal anti-human KLF4 antibody (H00009314-M01), chicken polyclonal anti-rabbit Immunoglobulin G (IgG) TRITC (NBP1-75270), and chicken polyclonal anti-mouse Immunoglobulin G (IgG) DyLight 594 (NBP1-75563) was purchased from Novus Biologicals (Littleton, CO). Goat anti-mouse immunoglobulin G (IgG)-horseradish peroxidase (HRP) (cat#170-6516) and goat anti-rabbit IgG-HRP (170-6515) were purchased from Bio-Rad (Hercules, CA, USA). Donkey anti-rabbit Alexa Flour-594 (A21207), donkey anti-mouse Alexa Flour-488 (A21202), TRIzol (15596018), and Lipofectamine™ 2000 (11668-019) were purchased from Invitrogen (Carlsbad, CA). 30% Acrylamide/Bis Solution, 29:1 (1610156) and Tetramethylethylenediamine (TEMED, 161-0801) were purchased from Bio-Rad (Hercules, CA, USA). Tamoxifen (J6350903, Alfa Aesar, Ward Hill, MA; T5648-1G, Sigma) stock was dissolved in corn oil (C8267, Sigma), filtered with a sterile 0.22μM filter, and stored at −20°C in a dark sealed container. It was warmed to room temperature prior to use. Lentivirus encoding control-shRNA and KLF4-shRNA knockdown vectors were purchased from Open Biosystems, Inc (RHS3979-201737586, RHS3979-201737587, RHS3979-201737588, RHS3979-201737589, RHS3979-201742512, Huntsville, AL, USA).

### Cell Culture and Media

These methods have been previously described (47-49,52,53). Primary human lung microvascular endothelial cells (hLMVECs) (CC-2527) were purchased from Lonza (Alpharetta, GA). Human umbilical vein endothelial cells (HUVECs) (SCCE001) were purchased from Millipore (Billerica, MA, USA). All primary endothelial cells (ECs) were cultured up to 3-5 passages in EndoGro^TM^ Basal Medium with EndoGro^TM^ VEGF Supplement Kit (SCME-BM; SCME002-S; millipore) on 0.2% gelatin (G-2625, Sigma-Aldrich) in PBS-coated plates. Amphopack 293T virus packaging cells and murine RAW 264.7 macrophage cells (gifted by Kostandin Pajcini, UIC) were cultured in Dulbecco’s minimum essential media (DMEM) (01-055-1, Biological Industries) supplemented with L-glutamine (25-005-CI; Corning) and 10% fetal bovine serum (F2442; Sigma-Aldrich).

### Co-Immunoprecipitation (Co-IP) Experiments

For Co-IP, adherent primary hLMVECs at 100% confluency were washed three times with cold 1x PBS, pH 7.4, followed by an incubation in 1x RIPA buffer on ice as previously described (47-49,52,53). For co-IP experiments, mouse IgG agarose beads and Protein-G Sepharose beads were washed three times with lysis buffer, and subsequently resuspended in PBS each to make 50% IgG agarose beads and 50% Protein-G Sepharose bead solutions, respectively. To reduce the amount of proteins that bind nonspecifically, the cell lysates were incubated with 30 μL of mouse IgG agarose beads on a rotator for 1 hour at 4°C. Beads/slurry were centrifuged at 5,000g for 30 seconds at 4°C and supernatant protein lysate transferred to a new 1.5 mL centrifuge tube. Immunocomplexes were washed 5 times with 1XTris, pH 7.5 buffer. Thereafter, each sample was diluted with 2X reducing sample buffer containing (5% β-mercaptoethanol) and boiled in a water bath for 5 minutes, and resolved by 10% SDS-PAGE, then transferred to as previously described (47-49,52-56).

### Human lung disease tissues and immunohistochemistry (IHC), and microscopy

The Lung Tissue Research Consortium (LTRC) at the National Health Institutes (NIH) has a collection of lung disease tissues. The program enrolled donors who underwent lung surgery, and blood and extensive phenotypic data were collected from the prospective donors, thereafter the surgical waste tissues are processed for research use. Most donor subjects harbored interstitial fibrotic lung disease, emphysema or COPD. Phenotypic data include clinical and pathological diagnoses, chest computed tomography (CT) scanning images, pulmonary function tests, exposure and symptom questionnaires, and exercise tests. Tissue samples used in this study were obtained from the LTRC for limited research use for biochemical and histological analyses. Portions of lung tissue samples were fixed in zinc formalin (10%) solution and embedded in paraffin as previously described (52–54). At least twelve to fifteen serial sections (6-8 µm thin) were prepared of each paraffin embedded organ. After deparaffinization, dehydration, and rehydration, the sections were permeabilized in 0.5% Triton X-100 in 1x PBS for 30 minutes and washed in 1X PBS. The sections were washed 3 times for 5 minutes in 1x PBS and incubated with secondary antibody (1:200, 2.5% BSA in 1x PBS) for 1 hr at RT in a humidified chamber, washed 3 times in 1x PBS. The slides were mounted using DAPI anti-fade gold reagent. Thereafter sections were examined under the Olympus BX51/IX70 fluorescence microscope (Olympus, Tokyo, Japan) and quantified using National Institute Health (NIH)-ImageJ software (Bethesda, MD, USA).

ECs up to passage 5 were seeded at 50-60% confluency on sterile glass coverslips coated with 0.02% gelatin in 1x PBS, pH 7.4. ECs were cultured for 4-5 days as described previously (47-49,52-55). Cells fixed in 4% PFA, washed in 1XPBS, permeabilized in 0.5% Triton X-100 (Fischer Scientific) in 1x PBS for 30 minutes at room temperature, followed by 3 washings in 1x PBS. Coverslips were incubated in 0.5% BSA, thereafter primary antibody applied for 1 hour (1:200 dilution with 0.5% BSA). After 1x PBS wash, secondary antibody (1:500 dilution with 0.5% BSA) was applied for 30 minutes. Coverslips were washed in 1x PBS and then mounted with ProLong® Diamond Antifade Mounting reagent with DAPI (P36962, Invitrogen). Digital microscopic images were acquired using Olympus BX51/IX70 Fluorescence microscope, the EVOS FL Cell Imaging System (InVitrogen), or Zeiss LSM 880 Confocal microscope saved as TIFF and CZI. Multiple panel figures were assembled using QuarkXpress 10.0 software. Final images were saves as EPS or TIFF documents.

### Production of pLentivirus and shRNA-KLF4 Knockdown

To amplify DNAs, we transformed *E. coli* (DH5α) with the five pLentivirus encoding shRNAs targeting *KLF4* gene and a scrambled shRNA control construct as previously described (47-49,52,53). To generate viral particles encoding the shRNAs, amphopack-293T cells were transfected with lentivirus encoding shRNA constructs using Lipofectamine as described previously (47,48). After 24 hours, transfected cells were fed with fresh media. After 36-48 hours, cells were detached and replated into 1:3 ratio in media containing 7 mg/ml puromycin (A11138-03, ThermoFischer Scientific). Puromycin media was changed every 3-4 days to remove dead cells. After 12-14 days, surviving cells/colonies were pooled and replated onto dishes. These cells were expanded into 1:3 ratio. After 3 days, one dish of cells was frozen, and two dishes were passaged for virus supernatant generation. Viral particles were collected in complete EC media and used immediately. The efficiency of knockdown was determined by RT-PCR and Western blot as previously described (47–49).

### Macrophage Cell Adhesion Assay

Cell adhesion assay has been previously described (47). ECs were grown to 60-80 percent confluency on 6 well plates coated with 0.02% gelatin in 1x PBS, pH 7.4 as previously described. ECs were treated with either control or KLF4 stealth siRNA for 6 hours, and 24 hours later were treated with either PBS or LPS (20 ug/mL) for 6 hrs. RAW 264.7 macrophages were labeled with mCherry using CellTracker™ Red CMTPX (C34552, Invitrogen) protocol. Fluorescently labeled RAW 264.7 macrophages were allowed to adhere to ECs for 0.5 hrs as previously described (47). ECs were washed to remove non-adherent cells and macrophage adherence was quantified by counting the number of fluorescent red macrophages per field using either the LSM 880 confocal or the EVOS FL Cell Imaging System.

### Quantitative (q)-reverse transcribed (RT)-PCR

To isolate total RNA, cells were grown to confluency and RNA samples were collected using either TRIzol or RNAzol RT reagent as previously described (54–56). The mRNAs were resuspended in 100 μl of RNAse free water and incubated at 55°C for 10 minutes, quantified, and stored at −80°C or used immediately. Automated qRT-PCR assay was carried out by a QuantStudio 7 Flex (Applied Biosystems) using SYBR green master mix (4367659, Applied Biosystems) with forward and reverse primers (Table 2) custom made through IDT (Coralville, IO). Data was analyzed using Δ-Δ-CT values normalized to housekeeping genes such as 18S-rRNA or GAPDH using Expression-Suite software (ThermoFisher Scientific).

### Generation of Klf4 genetically engineered mouse model (GEMM)

Mice were housed in a pathogen-free vivarium and handled in accordance with standard use procedures, animal welfare regulations, and the National Institutes of Health (NIH) Guide for the Care and Use of Laboratory Animals. All experimental procedures and husbandry have been approved by the University of Illinois at Chicago (UIC) Institutional Animal Care and Use Committee (IACUC). We procured Klf4^fl/wt^ mice on a C57BL background (Dr. Klaus Kaestner, University of Pennsylvania, Philadelphia, PA) (57), obtained from Mutant Mouse Resource & Research Centers (MMRRC) through an institutional Material Transfer Agreement (MTA). Introns 1 and 3 of the *Klf4* gene have *loxP* sites to create the *floxed* allele. These mice were crossbred to produce a generation of offspring such that a percentage of these mice were homozygous for the *Klf4^fl/fl^* mutant genotype. Our study used both male and female mice ranging from 12-14 weeks of age. For mouse experiments, no randomization was done, or no animals were excluded from this study. *Gt(Rosa)^26Sortm4(ACTB-tdTomato,-EGFP)Luo/J^* (stock#007576, hereafter called *Rosa^mT/mG^*) mice were bought from the Jackson Laboratory (Bar Harbor, ME) (58) and the *tg.Cdh5(PAC)^CreERT2^* (also known as *Tg:VE-cadherin-(PAC)^CreERT2^*, and hereafter called *Cdh5^CreERT2^*) transgenic mice were obtained from Ralph H. Adams (59) through an MTA from the UK Cancer Research (London, UK). Following TAM (2.0mg/kg body weight, BW) administration for each of five consecutive days, all ECs (expressing EC-specific *Cdh5^CreERT2^*) turn green due to mG expression, while non-ECs fluoresce tomato (red) due to baseline expression of mT. The transgenic *Cdh5^CreERT2^* driver line served as a proxy for EC specificity (53,57). After a series of breeding and backcrossing regimens, *Klf4^fl/fl^::Cdh5^CreERT2^*, *Rosa^mT/mG^::Cdh5^CreERT2^* and *Rosa^mT/mG^::Klf4^fl/fl^::Cdh5^CreERT2^* GEMM were produced.

### Genotyping of GEMM

Mouse tails were snipped (2mm) and the ears tagged aseptically as previously described (53,55). DNAs were isolated using Phenol:Chloroform:Isoamyl alcohol method (53,54). To induce deletion of EC-*Klf4* in a timed-manner, we injected 12-14 weeks old adult mice with 2.0mg/kg body weight TAM once every day intraperitoneally (i.p.) for 5 consecutive days. After treatment with TAM, we waited for 21 and 28 days so that all TAM were metabolized and excreted, thereafter mice were humanely sacrificed, different organs harvested, fixed, sectioned, and remaining tissues were processed for EC isolation, RNA and DNA extractions, and protein lysate preparation as previously described (53–55).

### Statistics

Data were calculated and expressed as mean ± S.D. and values plotted with Microsoft Excel or using GraphPad Prism 6 (GraphPad Software, La Jolla, CA, USA). For comparisons between two groups, an unpaired Student’s t-test was performed, and analysis of variance (ANOVA), followed by Tukey’s, Sidak’s, or Dunnett’s test for multiple comparisons. Number of experiments or number of animals represents biological replicates and a P value of < 0.05 was considered significant. qRT-PCR data expression was calculated using ΔΔCT values normalized against both 18s RNA and GAPDH of control samples.

## Supporting information

Supplemental Infor

## Authors Contributions

VM designed and executed experiments including immunohistology and microscopy, collected and analyzed data, generated graphics, and wrote the manuscript. CA was involved in animal husbandry, genotyping, and microscopy. AW conducted KLF4 and KLF2 amino acid sequence analysis, 2D and 3D-computational I-TASSER analysis. RDM advised and collaborated, read the manuscript, and provided feedbacks. KW designed experiments, supervised the project, and co-wrote the manuscript. All authors read and approved the final manuscript.

## Acknowledgments

The Research Resources Center (RRC) Core Microscopy and Imaging, Flow Cytometry Service, Research Histology, Cardiovascular Physiology Core, and Tissue Imaging Cores of the UIC.

## Supplemental Information

### Results

#### Computational analysis of KLF4(91-117aa) 2D- and 3D-structures

Amino acid residues 91-117 derived from KLF4-TAD were subjected to computational analysis via the I-TASSER online portal (58,59). Complete dataset is provided as a PDF document (Fig. S4). Predictive analysis shows that amino acid residues from KLF4(91-117) contains two discrete α-helical and two short stretches of coiled-coil regions as shown in Fig. 3F-G. Homology blast algorithm search identified Eukaryotic translation initiation factor 4E-binding protein (4E-BP)1 secondary structure as a mirror image of KLF4(91-117aa) secondary structure. As 4E-BP1 can bind several different substrates, we posit that KLF4(91-117) region is involved in the formation of homodimeric and heterodimeric protein complexes, such as KLF4-KLF4 and KLF4-KLF2 protein complexes. However, detailed analyses of these possibilities are beyond the scope of this report.

#### Generation of Rosa^mT/mG^::Klf4^fl/fl^::Cdh5^CreERT2^ mice

To address the *in vivo* role of KLF4 in EC quiescence, we prepared GEMM incorporating a tamoxifen-(TAM) inducible EC-specific deletion of *Klf4* alleles in a *Rosa^mT/mG^* reporter and C57BL (C57) genetic background (see methods) (Fig. S5). After several generations of breeding, we transferred *Rosa^mT/mG^* (dual-fluorescent reporter allele *mT/mG*) to *Cdh5^CreERT2^* TAM inducible mice in C57 background to generate *Rosa^mT/mG^::Cdh5^CreERT2^* mice. Thereafter, *Rosa^mT/mG^::Cdh5^CreERT2^* female mice were crossed with male *Klf4^fl/wt^* mice to produce heterozygous *Rosa^mT/mG^::Cdh5^CreERT2^::Klf4^fl/wt^* (50%). Next, we bred male *Rosa^mT/mG^::Cdh5^CreERT2^::Klf4^fl/wt^* with female *Rosa^mT/mG^::Cdh5^CreERT2^::Klf4^fl/wt^* to produce *Rosa^mT/mG^::Cdh5^CreERT2^::Klf4^fl/fl^* genotype (25%). For all downstream experiments, we used (1) *Rosa^mT/mG^::Klf4^fl/fl^* and (2) *Rosa^mT/mG^::Klf4^fl/fl^::Cdh5^CreERT2^* GEMM in C57BL background (Fig. S5A-C) and Cre-mediated recombination were confirmed by PCR genotyping (Fig. S5D).

### Supplemental Materials and Methods

#### Western Blotting (WB) analyses

WB methods have been previously described by us (49,52,53). Adherent 100% confluent primary hLMVECs were washed three times with 1x phosphate buffered saline (PBS), pH 7.4, followed by an incubation in 1x radioimmunoprecipitation assay (RIPA) buffer [50 mM 4-(2-hydroxyethel)-1-piperazineethanesulfonic acid (HEPES) (pH 7.4), 150 mM sodium chloride, 2 mM ethylenediaminetetraacetic acid (EDTA), 1.0% Triton X-100, 0.25% deoxycholic acid, 0.1% SDS, 10 mM tetrasodium pyrophosphate, 100 mM sodium fluoride, 0.98 mM sodium orthovanadate, and freshly added protease inhibitor (P8564, Sigma-Aldrich)] on ice for 5 minutes. For KLF4 and KLF2 oligomerization experiments, 100% confluent adherent hLMVECs were washed three times with 1x PBS, pH 7.4, followed by an incubation in 1x Tris cell extraction buffer [20 mM Tris (pH 7.4), 150 mM NaCl, 2 mM EDTA (pH 8.0), 0.5% Triton X-100, 10 mM tetrasodium pyrophosphate, 100 mM sodium fluoride, 0.98 mM sodium orthovanadate, and freshly added protease inhibitor] on ice for 5 minutes. Protein extracts were clarified by centrifugation at 21,000g (14,000rpm) at 4°C for 20 minutes, supernatant saved, and total protein concentration quantified. Each sample was diluted with 2x reducing sample buffer [50 mM Tris, 2.0% SDS, 10% glycerol, 0.01% bromophenol blue, 5% β-mercaptoethanol, pH 6.8] and boiled in water bath for 5 minutes. Proteins complexes were resolved by 8% sodium dodecyl sulfate (SDS)-polyacrylamide gel electrophoresis (PAGE) and ECL signals were developed by an X-ray developer and signals quantified using ImageJ as previously described (47-49,52,53).

#### Chromatin Immunoprecipitation (ChIP) and Electrophoretic Mobility Shift Assays (EMSA)

For ChIP experiments, hLMVECs were grown to 100% confluency on plates coated with 0.02% gelatin in 1x PBS, pH 7.4 as described above. Each plate of cells was incubated at 37°C for 10 minutes in 1.0% formaldehyde (Thermo-Fisher, Rockford, IL), thereafter, reverse crosslinked, washed, processed and DNAs sheared. Chromatins were pre-cleared, immunoprecipitated with anti-IgG, anti-KLF2, or anti-KLF4 antibodies as previously described (49,52). For EMSA, nuclear extracts from hLMVECs were prepared according to the manufacturer’s instructions (Thermo-Fisher) as previously described (47-49,53). For EMSA, KLF4-binding oligonucleotides modeled after the KLF2-promoter were synthesized by (IDT DNA Corporation) and labeled with biotin. Protein-DNA complexes were analyzed by 10% native PAGE. Thereafter, nitrocellulose (NC) membranes were subjected to streptavidin-horseradish peroxidase (HRP) and enhanced chemiluminescence (ECL) analyses.

#### Isolation of mouse lung microvessel endothelial cells (mLMVECs)

Microbeads coated with anti-mouse CD31 (Miltenyi Biotech., 130-097-418) and anti-mouse CD45 antibodies (Miltenyi Biotech., 130-052-301) were used to isolate ECs from mouse tissue as previously described (53–57). For EC isolation, mice were humanely sacrificed. Lungs from 2 mice were combined in a 10 cm dish filled with ice cold DMEM media and cut into fine pieces with sterile scissors. Tissue was transferred to a 15 mL conical tube filled with 12 mL of digestion solution [1 mg/mL collagenase IV, 10 mg/mL BSA, 1 U/mL Dispase in DMEM] and incubated at 37°C for 45 minutes on a rotator, suspended with a 14 gauge cannula every four minutes to get even cell suspension. Cell suspension was passed through a 70 µm cell strainer and washed with 15 mL of plain DMEM media to halt digestion. The cell suspension was centrifuged at 400xg for 5 minutes and then supernatant was aspirated. The cell pellet was resuspended in 3 mL of sterile 0.22 µm filtered 0.1% BSA in 1x PBS and transferred to 5 mL round bottom polystyrene tubes. Next, cell suspension was incubated with 22.5 μL of the prepared anti-mouse CD45 antibody at 4°C for 30 minutes. CD45+ cells discarded, while anti-CD45 negative cells were mixed with anti-mouse CD31 antibody conjugated Microbeads at 4°C for 30 minutes on an end over end rocker. Polystyrene tubes were exposed to the magnet again. To release the cells from the beads, they were incubated in 5mM EDTA on a rotator at 37°C for 10 minutes. The tube was then placed on the magnet, and supernatant containing the isolated EC population was collected and then used for downstream experiments.

#### Hematoxylin and Eosin (H&E) and Trichrome Staining, and Microscopy

Mice were anesthetized and sacrificed humanely as previously described (53–57). For cryosections, lungs were perfused with PBS, collected, rinsed in PBS, and placed in increasing concentrations of PBS/sucrose solution overnight at 4°C (3%, 10%, 30%) for cryoprotection and in sucrose:OCT (TissueTek 4583) (30% in PBS) until the specimen reached the bottom of the mold. These samples were frozen in OCT and stored at −80°C. Cryosections were sectioned at 6-8 μm thickness using a Cryostat. Sections were then washed in distilled water, followed by PBS three times. Slides were mounted using prolong anti-FADE mounting solution containing DAPI (InVitrogen). Fluorescence images were recorded using Olympus microscope at room temperature.

For H&E and Trichrome staining, freshly isolated lung tissues were washed in cold PBS and fixed immediately in zinc formalin (10%) overnight at 4°C and embedded in paraffin. At least twelve to fifteen serial sections (6-8 μm thin) were prepared. After deparaffinization, sections were permeabilized in 0.5% Triton X-100, washed in 1XPBS, incubated in citrate buffer (10 mM citric acid, 0.05% Tween-20, pH 6.0, Tri-sodium citrate, 2.94g, distilled H_2_O 1L) at 90°C for an hour and a half in a water bath. After several washes, slides were briefly treated with hydrogen peroxide, blocked in 3% BSA prepared in 1x PBS for 30 minutes. Next, sections were incubated with primary antibody (1.0-2.0 μg/ml) for 1 hr at RT in a humidified chamber, washed 5 times, mounted using DAPI anti-fade gold reagent, Thereafter, examined under the Olympus BX51/IX70 fluorescence microscope (Olympus, Tokyo, Japan). Morphometric analyses of lung architecture were analyzed using National Institute Health (NIH)-ImageJ software (Bethesda, MD, USA) as previously described (53–56). In brief, TIFF images analyzed by NIH-ImageJ, where a command was run to subtract background from all images, and to separate red and blue staining from each other color deconvolution was performed. Each image was subsequently quantified for red and blue staining. Blue staining correlated with collagen deposition; using a ratio of blue quantification to red quantification for each field, collagen staining could be normalized to the red staining of the general architecture. At least 5 fields each in 5 mice per experimental group were quantified for each organ.

### Supplemental Figure legends

**Figure S1. *KLF4* and *KLF2* transcripts in human tissues. A&B.** Screen grabs of RNA-seq analyses show *Klf4* (www.ncbi.nlm.nih.gov/gene/9314) and *KLF2* (www.ncbi.nlm.nih.gov/gene/10365) transcripts in normal human tissues, and abundantly found in the lung.

**Figure S2. Analysis of KLF4, KLF2, and TGF-β1 in human lung inflammatory disease samples. A&B**. Control healthy donors **C&D**. Lung tissues obtained from emphysema and COPD patients were stained with anti-KLF2 or KLF4. Representative photomicrographs of remodeled pulmonary vessels, decreased alveolar counts in COPD patients, compared with healthy vessels in donors are shown. **E**. Quantification of KLF2 and KLF4 positive nuclei (dark brown). Data represents mean ± S.D. and p values and “n” are as shown. **F**. The cell lysates were subjected to Western blotting (WB) with antibodies against KLF4, KLF2, TGF-β1 and GAPDH. GAPDH was used as a control for equal protein loading. The numbers below each panel represent signal intensities relative to GAPDH. Experiments were repeated at three times. n = number of samples. P<0.05 is considered statistically significant. Scale bar and magnifications areas are as shown.

**Figure S3. Human lung disease harbor microscopic EndMT.** A thin section (6-7μ) of emphysema tissue section was immunostained with anti-CD31 (green) and anti-α-SMA (red) antibodies and analyzed by epifluorescence microscopy at indicated magnification. Blue color indicates nuclear staining (DAPI). Magnifications and scale bars are as shown. The brightness and contrast of this image was not adjusted. Quantification was not performed due to limited sample size.

**Figure S4. KLF4(91-117aa) 2D and 3D structure computational analysis.** Amino acid residues -NDLLDLDFILSNSLTHPPESVAATVS-was submitted for Iterative Threading ASSEmbly Refinement (I-TASSER) (https://zhanglab.dcmb.med.umich.edu/I-TASSER/). I-TASSER is a protein structure prediction and structure-based function annotation tool. It identifies and retrieves structural templates from the Protein Database (PDB) by multiple threading method LOMETS, with full-length atomic models constructed by iterative template-based fragment assembly simulations.

**Figure S5. Generation of *Rosa^mT/mG^::Klf4^fl/fl^::Cdh5^CreERT2^* mice line. A**. The timeline and TAM treatment scheme; **E**. Representative ethidium-bromide stained agarose gel images of PCR genotyping. Genomic DNA of *wild-type (WT) Klf4, floxed (fl) Klf4, Cdh5^CreERT2^ (Cre), and Rosa^mT/mG^* mice were amplified by PCR. The WT band is 464 bp, the floxed Klf4 product is 618 bp, and the *Cdh5^CreERT2^* product is 934 bp. *MW, ladder*. **F**. EC extracts were prepared from indicated mouse lung tissues and analyzed by WB with indicated antibodies. Molecular weights are given in kiloDalton (kDa). The numbers below each WB panel indicate signal quantification. Experiments were performed at least 3 times.

