## Supplemental Infor for "A requirement for Krüppel-Like Factor-4 in the maintenance of endothelial cell quiescence"

### Fig. S1

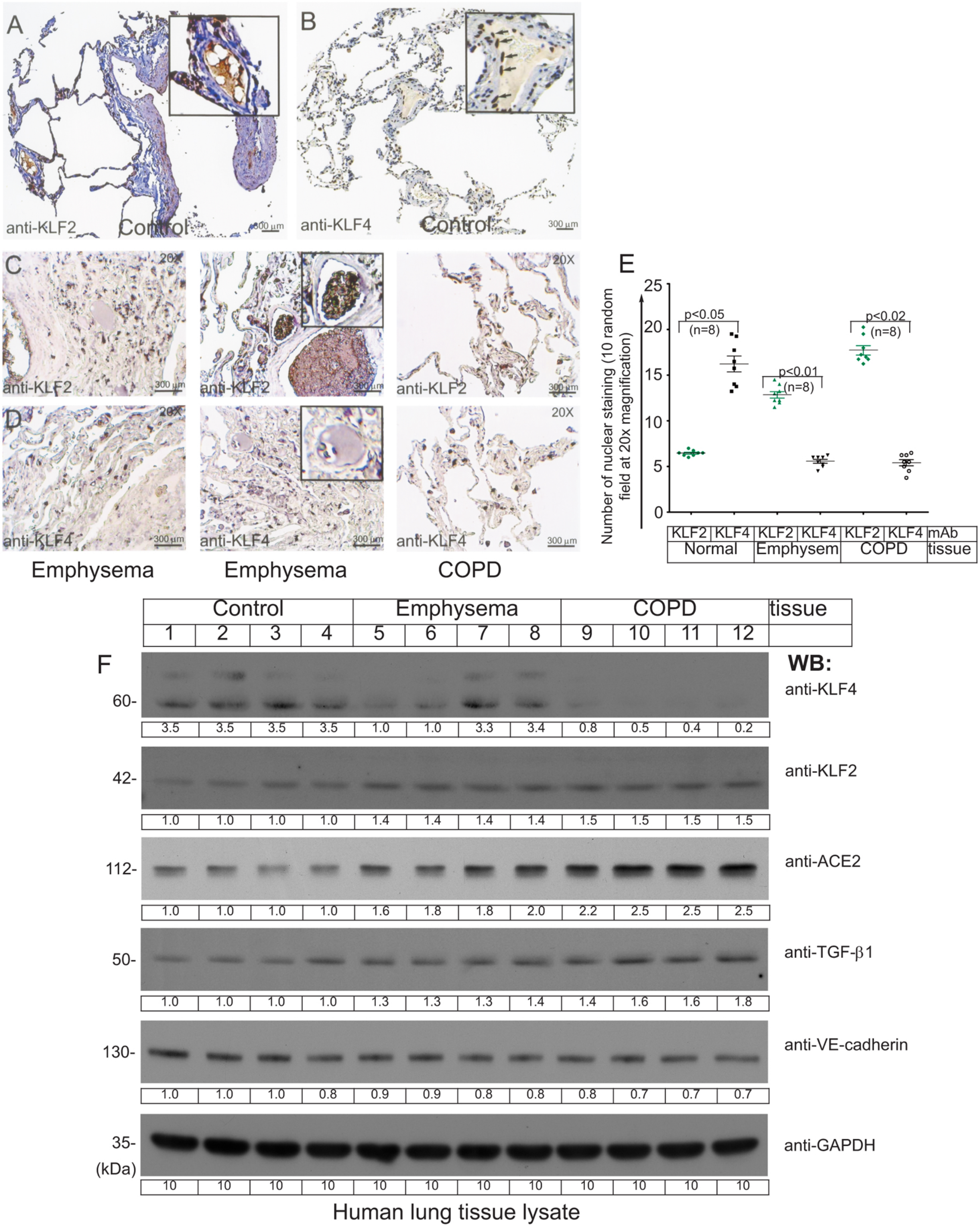

A
anti-KLF2
Control
300 µm
B
anti-KLF4
Control
300 µm
C
20X
anti-KLF2
anti-KLF2
anti-KLF2
D
anti-KLF4
anti-KLF4
anti-KLF4
Emphysema
Emphysema
COPD
E
Number of nuclear staining (10 random field at 20x magnification)
p<0.05 (n=8)
p<0.01 (n=8)
p<0.02 (n=8)
KLF2 KLF4 KLF2 KLF4 KLF2 KLF4 mAb
Normal Emphysem COPD tissue
Control
Emphysema
COPD
tissue
1 2 3 4 5 6 7 8 9 10 11 12
F
WB:
60- anti-KLF4
3.5 3.5 3.5 3.5 1.0 1.0 3.3 3.4 0.8 0.5 0.4 0.2
42- anti-KLF2
1.0 1.0 1.0 1.0 1.4 1.4 1.4 1.4 1.5 1.5 1.5 1.5
112- anti-ACE2
1.0 1.0 1.0 1.0 1.6 1.8 1.8 2.0 2.2 2.5 2.5 2.5
50- anti-TGF-β1
1.0 1.0 1.0 1.0 1.3 1.3 1.3 1.4 1.4 1.6 1.6 1.8
130- anti-VE-cadherin
1.0 1.0 1.0 0.8 0.9 0.9 0.8 0.8 0.8 0.7 0.7 0.7
35- anti-GAPDH
(kDa)
10 10 10 10 10 10 10 10 10 10 10 10
Human lung tissue lysate

Fig. S2

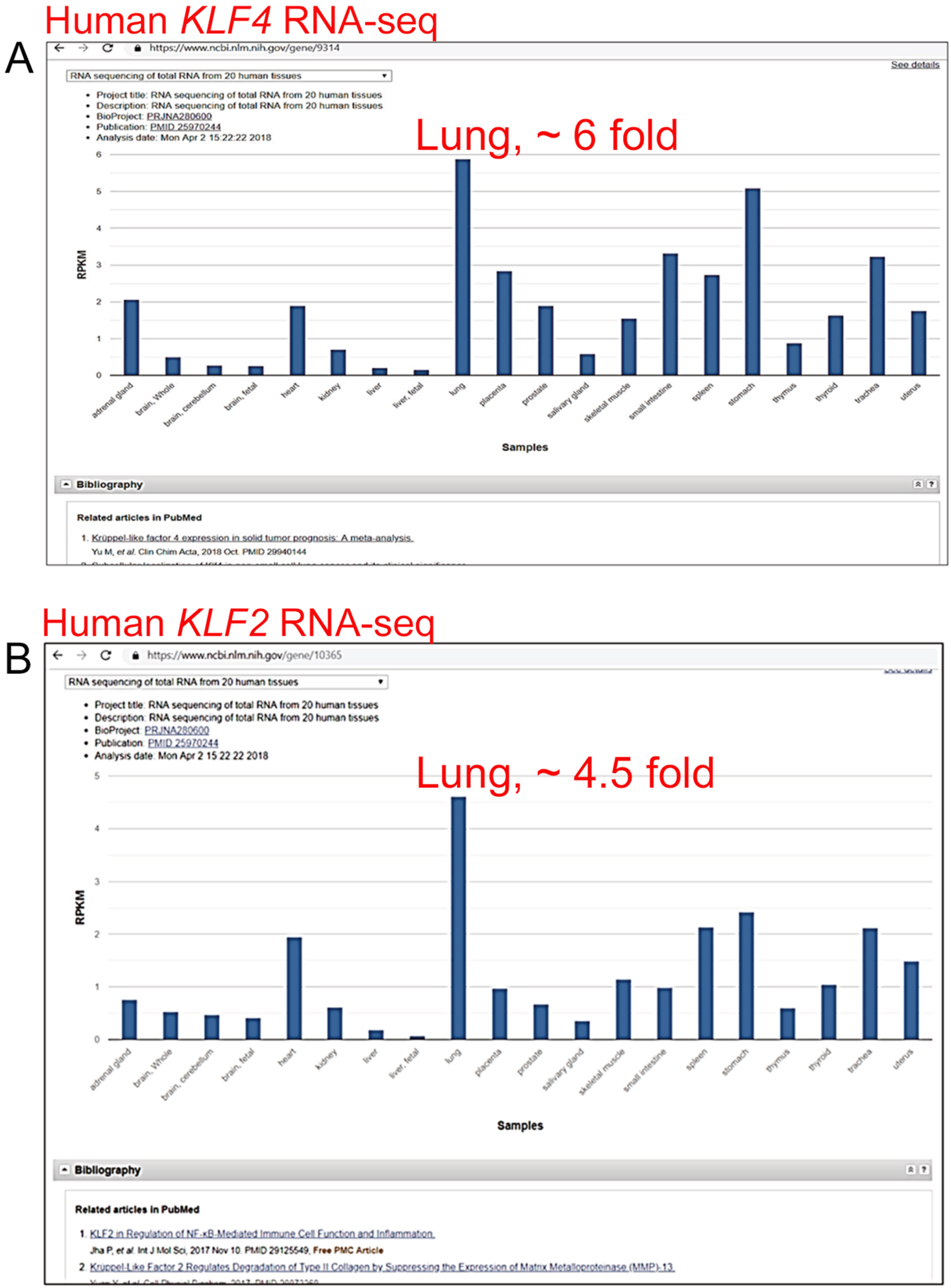
Human *KLF4* RNA-seq
A
https://www.ncbi.nlm.nih.gov/gene/9314
See details
RNA sequencing of total RNA from 20 human tissues
Project title: RNA sequencing of total RNA from 20 human tissues
Description: RNA sequencing of total RNA from 20 human tissues
BioProject: PRJNA280600
Publication: PMID 25970244
Analysis date: Mon Apr 2 15:22:22 2018
Lung, ~ 6 fold
RPKM
0
1
2
3
4
5
6
adrenal gland
brain, Whole
brain, cerebellum
brain, fetal
heart
kidney
liver
liver, fetal
lung
placenta
prostate
salivary gland
skeletal muscle
small intestine
spleen
stomach
thymus
thyroid
trachea
uterus
Samples
Bibliography
Related articles in PubMed
1. Krüppel-like factor 4 expression in solid tumor prognosis: A meta-analysis.
Yu M, et al. Clin Chim Acta, 2018 Oct. PMID 29940144
Human *KLF2* RNA-seq
B
https://www.ncbi.nlm.nih.gov/gene/10365
RNA sequencing of total RNA from 20 human tissues
Project title: RNA sequencing of total RNA from 20 human tissues
Description: RNA sequencing of total RNA from 20 human tissues
BioProject: PRJNA280600
Publication: PMID 25970244
Analysis date: Mon Apr 2 15:22:22 2018
Lung, ~ 4.5 fold
RPKM
0
1
2
3
4
5
adrenal gland
brain, Whole
brain, cerebellum
brain, fetal
heart
kidney
liver
liver, fetal
lung
placenta
prostate
salivary gland
skeletal muscle
small intestine
spleen
stomach
thymus
thyroid
trachea
uterus
Samples
Bibliography
Related articles in PubMed
1. KLF2 in Regulation of NF-κB-Mediated Immune Cell Function and Inflammation.
Jha P, et al. Int J Mol Sci, 2017 Nov 10. PMID 29125549, Free PMC Article
2. Kruppel-Like Factor 2 Regulates Degradation of Type II Collagen by Suppressing the Expression of Matrix Metalloproteinase (MMP)-13.

Fig. S3

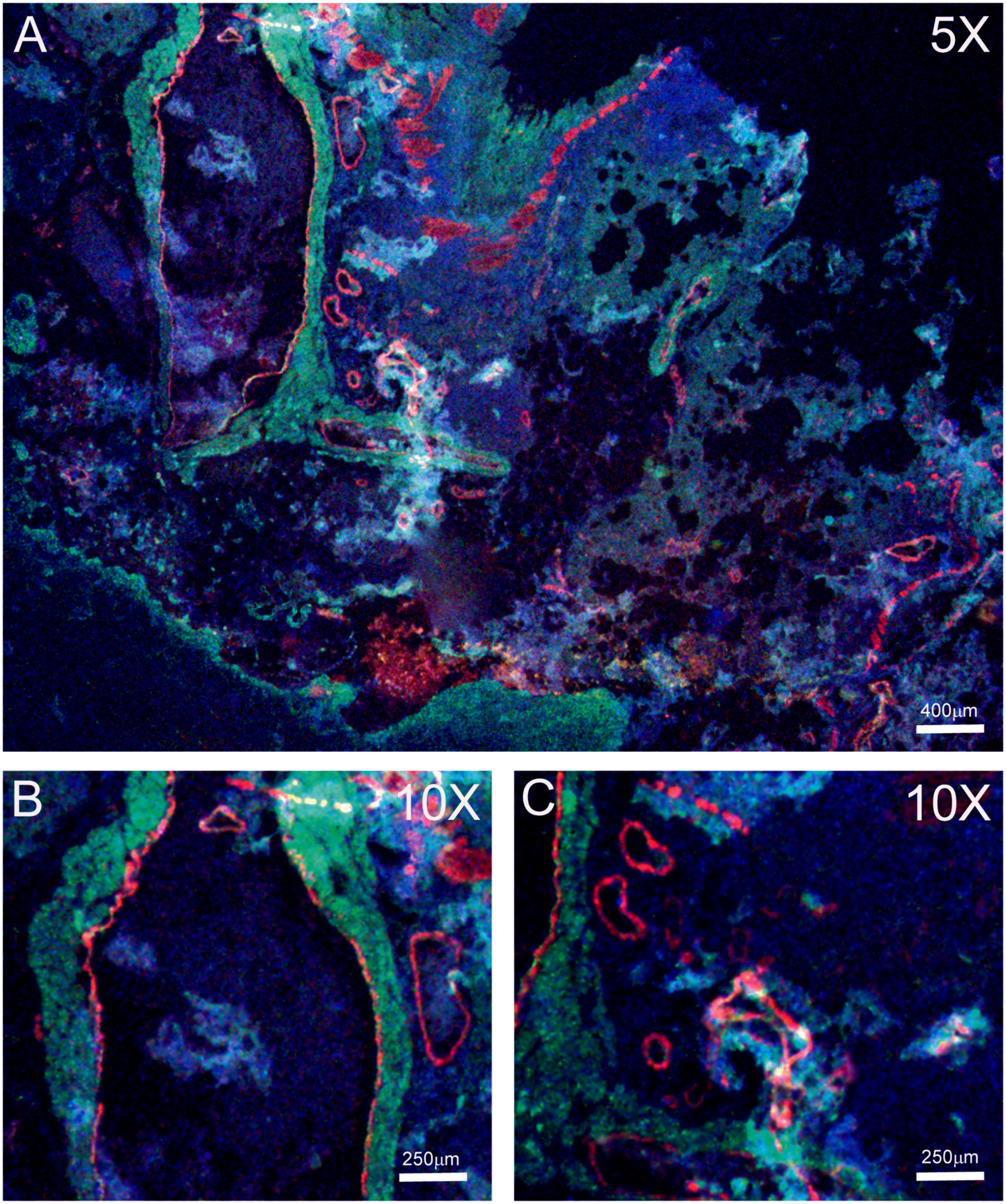

**Green:** CD31 **Red:** αSMA **Blue:** DAPI

Fig. S4

[Home] [Server] [Queue] [About] [Remove] [Statistics]

### I-TASSER results for job id S609161

(Click on S609161_results.tar.bz2 to download the tarball file including all modeling results listed on this page. Click on Annotation of I-TASSER Output to read the instructions for how to interpret the results on this page. Model results are kept on the server for 60 days, there is no way to retrieve the modeling data older than 2 months)

#### Submitted Sequence in FASTA format

```
>seq
NDLLDLDFILSNSLTHPPESVAATVS
```

#### Predicted Secondary Structure

```
                        20
                         |
Sequence   NDLLDLDFILSNSLTHPPESVAATVS
Prediction CCCHHHHHHHHCCCCCCCHHHHHCCC
Conf.Score 94122445521666798177630149
           H:Helix; S:Strand; C:Coil
```

#### Predicted Solvent Accessibility

```
                        20
                         |
Sequence   NDLLDLDFILSNSLTHPPESVAATVS
Prediction 84334143214442443464244438
           Values range from 0 (buried residue) to 9 (highly exposed residue)
```

#### Predicted normalized B-factor

(B-factor is a value to indicate the extent of the inherent thermal mobility of residues/atoms in proteins. In I-TASSER, this value is deduced from threading template proteins from the PDB in combination with the sequence profiles derived from sequence databases. The reported B-factor profile in the figure below corresponds to the normalized B-factor of the target protein, defined by B=(B'-u)/s, where B' is the raw B-factor value, u and s are respectively the mean and standard deviation of the raw B-factors along the sequence. Click here to read more about predicted normalized B-factor)

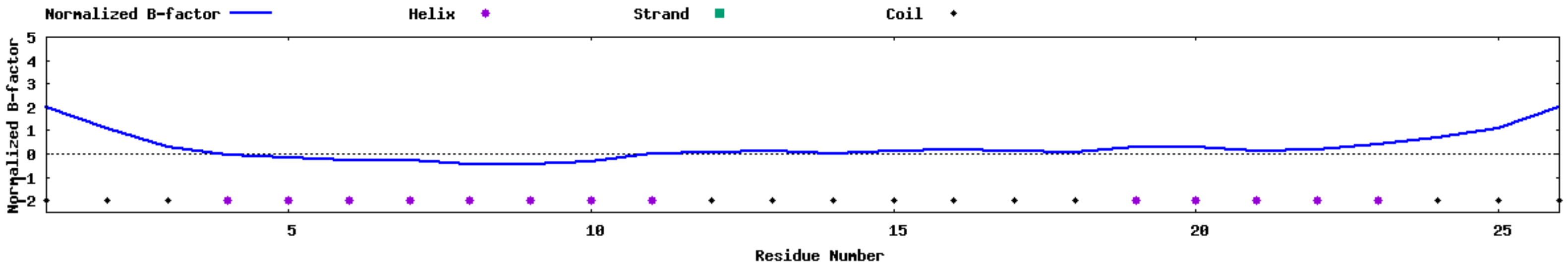

#### Top 10 threading templates used by I-TASSER

(I-TASSER modeling starts from the structure templates identified by LOMETS from the PDB library. LOMETS is a meta-server threading approach containing multiple threading programs, where each threading program can generate tens of thousands of template alignments. I-TASSER only uses the templates of the highest significance in the threading alignments, the significance of which are measured by the Z-score, i.e. the difference between the raw and average scores in the unit of standard deviation. The templates in this section are the 10 best templates selected from the LOMETS threading programs. Usually, one template of the highest Z-score is selected from each threading program, where the threading programs are sorted by the average performance in the large-scale benchmark test experiments.)

| Rank | PDB Hit | Iden1 | Iden2 | Cov | Norm. Z-score | Download Align. | 20<br>\| |
|---|---|---|---|---|---|---|---|

| | | | | | | | |
|---|---|---|---|---|---|---|---|
| | | | | | | **Sec.Str** | CCCHHHHHHHHCCCCCCCHHHHHCCC |
| | | | | | | **Seq** | NDLLDLDFILSNSLTHPPESVAATVS |
| 1 | 4uedB | 0.31 | 0.19 | 1.00 | 1.18 | Download | RIIYDRKFLMENSVTKPPRDLPTIVT |
| 2 | 5yfpC | 0.12 | 0.15 | 0.96 | 1.13 | Download | QYYKELHSLI-TDLVEPETIIILDIL |
| 3 | 5oqlJ | 0.15 | 0.15 | 1.00 | 1.27 | Download | LKLVEQEQTLENYIHAGAYRDAIVLA |
| 4 | 2auhB | 0.33 | 0.31 | 0.92 | 1.16 | Download | NSLVAMDF--SGQIENPTEALSVAVE |
| 5 | 7kchA | 0.04 | 0.19 | 0.88 | 1.12 | Download | VETVWIE---ATVVKLDGDAITARTV |
| 6 | 1g1pA | 0.27 | 0.27 | 1.00 | 1.13 | Download | DDCIKYGFILKNGLCCSGACVGVCAD |
| 7 | 4xfvA | 0.23 | 0.23 | 1.00 | 1.01 | Download | FTHERMDFFLTNGKTGSWRMWATCDQ |
| 8 | 5mqfF | 0.23 | 0.27 | 1.00 | 1.09 | Download | TGLSEGSYLLSNAMDNPKERCVKIFQ |
| 9 | 6c4lC | 0.17 | 0.19 | 0.92 | 1.00 | Download | --INLLRFLNDDNYPVPASMVSRRWI |
| 10 | 2waxB | 0.19 | 0.19 | 1.00 | 1.09 | Download | EEIPDTDFDFEGNLAFDKAAVFEEID |

(a) All the residues are colored in black; however, those residues in template which are identical to the residue in the query sequence are highlighted in color. Coloring scheme is based on the property of amino acids, where polar are brightly coloured while non-polar residues are colored in dark shade. (more about the colors used)

(b) Rank of templates represents the top ten threading templates used by I-TASSER.

(c) Ident1 is the percentage sequence identity of the templates in the threading aligned region with the query sequence.

(d) Ident2 is the percentage sequence identity of the whole template chains with query sequence.

(e) Cov represents the coverage of the threading alignment and is equal to the number of aligned residues divided by the length of query protein.

(f) Norm. Z-score is the normalized Z-score of the threading alignments. Alignment with a Normalized Z-score >1 mean a good alignment and vice versa.

(g) Download Align. provides the 3D structure of the aligned regions of the threading templates.

(h) The top 10 alignments reported above (in order of their ranking) are from the following threading programs:

1: MUSTER 2: SPARKS-X 3: Neff-PPAS 4: MUSTER 5: SPARKS-X 6: MUSTER 7: SPARKS-X 8: MUSTER 9: SPARKS-X 10: MUSTER

#### Top 5 final models predicted by I-TASSER

(For each target, I-TASSER simulations generate a large ensemble of structural conformations, called decoys. To select the final models, I-TASSER uses the SPICKER program to cluster all the decoys based on the pair-wise structure similarity, and reports up to five models which corresponds to the five largest structure clusters. The confidence of each model is quantitatively measured by C-score that is calculated based on the significance of threading template alignments and the convergence parameters of the structure assembly simulations. C-score is typically in the range of [-5, 2], where a C-score of a higher value signifies a model with a higher confidence and vice-versa. TM-score and RMSD are estimated based on C-score and protein length following the correlation observed between these qualities. Since the top 5 models are ranked by the cluster size, it is possible that the lower-rank models have a higher C-score in rare cases. Although the first model has a better quality in most cases, it is also possible that the lower-rank models have a better quality than the higher-rank models as seen in our benchmark tests. If the I-TASSER simulations converge, it is possible to have less than 5 clusters generated; this is usually an indication that the models have a good quality because of the converged simulations.)

- More about C-score
- Local structure accuracy profile of the top five models

(By right-click on the images, you can export image file or change the configurations, e.g. modifying the background color or stopping the spin of your models)

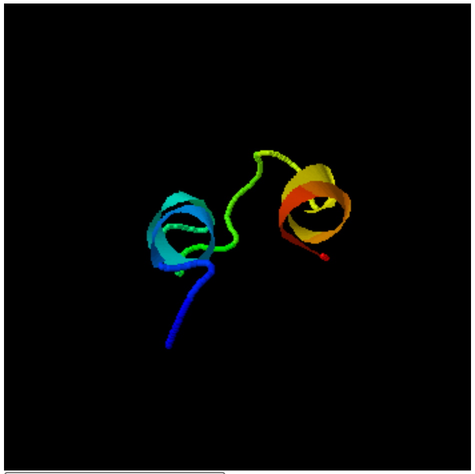

Reset to initial orientation Spin On/Off

- Download Model 1
- C-score=-1.37 (Read more about C-score)
- Estimated TM-score = 0.55±0.15

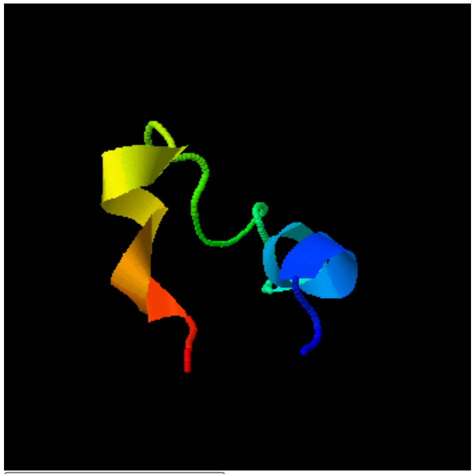

Reset to initial orientation Spin On/Off

- Download Model 2
- C-score = -2.53

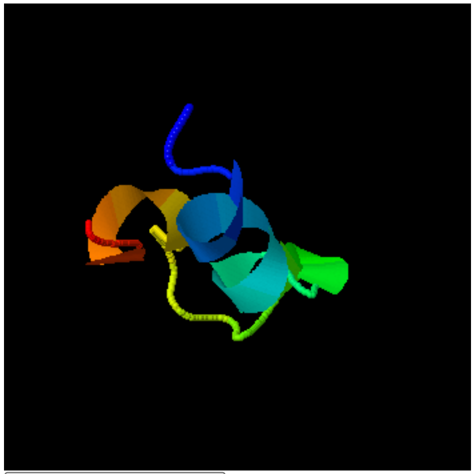

Reset to initial orientation Spin On/Off

- Download Model 3
- C-score = -1.95

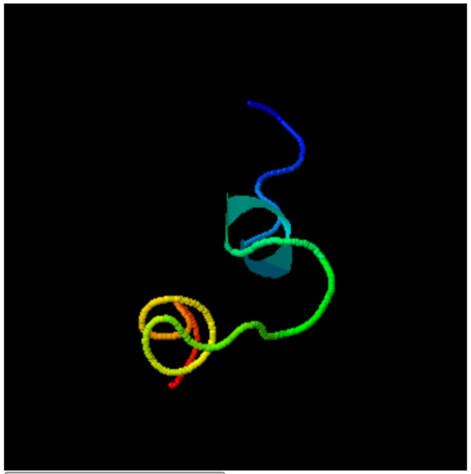

Reset to initial orientation Spin On/Off

- Download Model 4
- C-score = -3.55

- Estimated RMSD = 4.0±2.7Å

#### Proteins structurally close to the target in the PDB (as identified by TM-align)

(After the structure assembly simulation, I-TASSER uses the TM-align structural alignment program to match the first I-TASSER model to all structures in the PDB library. This section reports the top 10 proteins from the PDB that have the closest structural similarity, i.e. the highest TM-score, to the predicted I-TASSER model. Due to the structural similarity, these proteins often have similar function to the target. However, users are encouraged to use the data in the next section 'Predicted function using COACH' to infer the function of the target protein, since COACH has been extensively trained to derive biological functions from multi-source of sequence and structure features which has on average a higher accuracy than the function annotations derived only from the global structure comparison.)

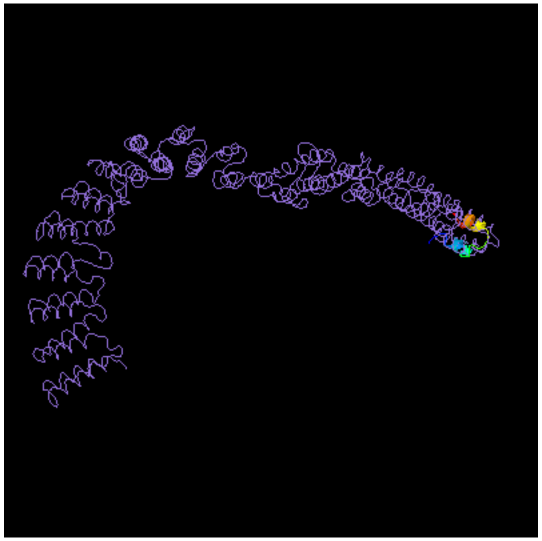

**Top 10 Identified stuctural analogs in PDB**

| Click to view | Rank | PDB Hit | TM-score | RMSD[a] | IDEN[a] | Cov | Alignment |
|---|---|---|---|---|---|---|---|
| ○ | 1 | 7abiM | 0.537 | 2.06 | 0.080 | 0.923 | Download |
| ○ | 2 | 3dm5A1 | 0.503 | 2.65 | 0.080 | 0.962 | Download |
| ○ | 3 | 2wviA1 | 0.493 | 2.22 | 0.154 | 0.962 | Download |
| ○ | 4 | 6w2vA2 | 0.489 | 1.99 | 0.000 | 0.923 | Download |
| ○ | 5 | 4uf1A | 0.483 | 2.16 | 0.125 | 0.923 | Download |
| ○ | 6 | 3oo6A2 | 0.482 | 2.14 | 0.174 | 0.885 | Download |
| ○ | 7 | 1hxgA | 0.478 | 1.97 | 0.160 | 0.923 | Download |
| ○ | 8 | 2d5mA | 0.476 | 2.56 | 0.042 | 0.923 | Download |
| ○ | 9 | 3tduA | 0.476 | 2.20 | 0.000 | 0.923 | Download |
| ○ | 10 | 6tgbA9 | 0.475 | 2.09 | 0.000 | 0.923 | Download |

(a) Query structure is shown in cartoon, while the structural analog is displayed using backbone trace.
(b) Ranking of proteins is based on TM-score of the structural alignment between the query structure and known structures in the PDB library.
(c) RMSD[a] is the RMSD between residues that are structurally aligned by TM-align.
(d) IDEN[a] is the percentage sequence identity in the structurally aligned region.
(e) Cov represents the coverage of the alignment by TM-align and is equal to the number of structurally aligned residues divided by length of the query protein.

Reset to initial orientation ☐ Spin On/Off

#### Predicted function using COFACTOR and COACH

(This section reports biological annotations of the target protein by COFACTOR and COACH based on the I-TASSER structure prediction. While COFACTOR deduces protein functions (ligand-binding sites, EC and GO) using structure comparison and protein-protein networks, COACH is a meta-server approach that combines multiple function annotation results (on ligand-binding sites) from the COFACTOR, TM-SITE and S-SITE programs.)

##### Ligand binding sites

| Click to view | Rank | C-score | Cluster size | PDB Hit | Lig Name | Download Complex | Ligand Binding Site Residues |
|---|---|---|---|---|---|---|---|
| ◉ | 1 | 0.21 | 15 | 2wksB | CB6 | Rep, Mult | 7,10,11 |
| ○ | 2 | 0.15 | 10 | 3tz3A | B36 | Rep, Mult | 5,6,8,9 |
| ○ | 3 | 0.13 | 9 | 3nwyA | GTP | Rep, Mult | 3,4 |
| ○ | 4 | 0.03 | 2 | 1g6uA | TFA | Rep, Mult | 20,21,22 |
| ○ | 5 | 0.01 | 1 | N/A | N/A | N/A | 8,9,10,11,15,17 |

Download the residue-specific ligand binding probability, which is estimated by SVM.
Download the all possible binding ligands and detailed prediction summary.
Download the templates clustering results.
(a) **C-score** is the confidence score of the prediction. C-score ranges [0-1], where a higher score indicates a more reliable prediction.
(b) **Cluster size** is the total number of templates in a cluster.
(c) **Lig Name** is name of possible binding ligand. Click the name to view its information in the BioLiP database.

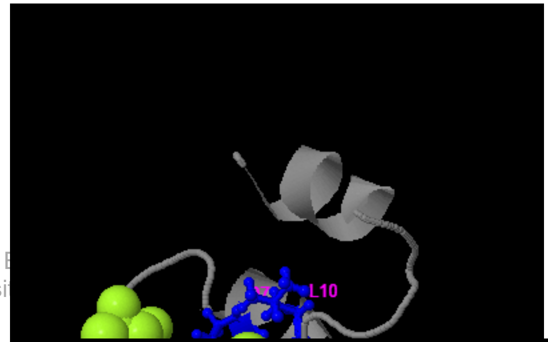

(d) **Rep** is a single complex structure with the most representative ligand in the cluster, i.e., the one listed in the **Lig Name** column. **Mult** is the complex structures with all potential binding ligands in the cluster.

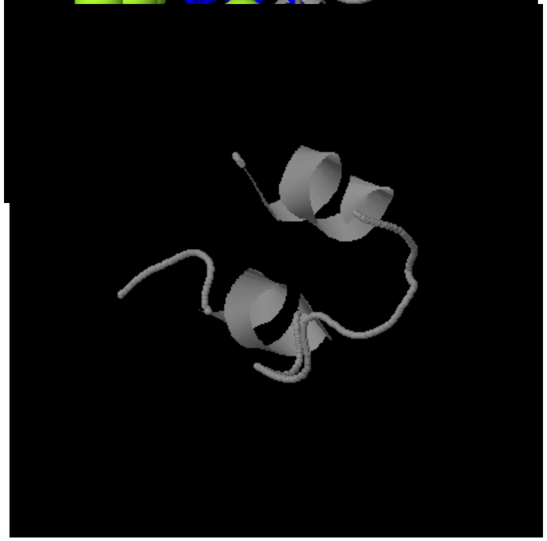

| Click to view | Rank | Cscore$^{EC}$ | PDB Hit | TM-score | RMSD$^a$ | IDEN$^a$ | Cov | EC Number | Active Site Residues |
|---|---|---|---|---|---|---|---|---|---|
| ◯ | 1 | 0.144 | 1wowA | 0.433 | 2.41 | 0.040 | 0.923 | 1.14.99.3 | NA |
| ◯ | 2 | 0.143 | 2ksfA | 0.432 | 2.59 | 0.000 | 0.885 | 2.7.13.3 | NA |
| ◯ | 3 | 0.142 | 1wcgA | 0.437 | 2.62 | 0.040 | 0.885 | 3.2.1.147 | NA |
| ◯ | 4 | 0.136 | 1xqoA | 0.439 | 2.75 | 0.000 | 0.923 | 4.2.99.18 | NA |
| ◯ | 5 | 0.136 | 2oqdB | 0.436 | 3.08 | 0.080 | 0.962 | 3.1.1.4 | NA |

**Click on the radio buttons to visualize predicted active site residues.**

(a) Cscore$^{EC}$ is the confidence score for the EC number prediction. Cscore$^{EC}$ values range in between [0-1]; where a higher score indicates a more reliable EC number prediction.

(b) TM-score is a measure of global structural similarity between query and template protein.

(c) RMSD$^a$ is the RMSD between residues that are structurally aligned by TM-align.

(d) IDEN$^a$ is the percentage sequence identity in the structurally aligned region.

(e) Cov represents the coverage of global structural alignment and is equal to the number of structurally aligned residues divided by length of the query protein.

Reset to initial orientation ☐ Spin On/Off

#### Gene Ontology (GO) terms

##### Top 10 homologous GO templates in PDB

| Rank | Cscore$^{GO}$ | TM-score | RMSD$^a$ | IDEN$^a$ | Cov | PDB Hit | Associated GO Terms |
|---|---|---|---|---|---|---|---|
| 1 | 0.17 | 0.3300 | 2.15 | 0.12 | 0.92 | 3oo9A | GO:0005215 GO:0006810 |
| 2 | 0.17 | 0.4882 | 2.29 | 0.08 | 0.88 | 3dm5A | GO:0008312 GO:0030529 GO:0006614 GO:0005737 GO:0017111 GO:0003723 GO:0005786 GO:0005525 GO:0000166 GO:0048500 |
| 3 | 0.16 | 0.4384 | 2.65 | 0.04 | 0.88 | 3q0oB | GO:0003723 GO:0005488 |
| 4 | 0.15 | 0.4503 | 2.51 | 0.19 | 1.00 | 1yhuC | GO:0005506 GO:0005576 GO:0005833 GO:0015671 GO:0019825 GO:0020037 |
| 5 | 0.14 | 0.3969 | 3.00 | 0.08 | 0.96 | 3h3dY | GO:0003723 GO:0005488 |
| 6 | 0.14 | 0.4144 | 2.83 | 0.00 | 0.92 | 3q0qA | GO:0003723 GO:0005488 |
| 7 | 0.13 | 0.4489 | 2.08 | 0.00 | 0.92 | 2yinA | GO:0005085 GO:0005506 GO:0005525 GO:0009055 GO:0020037 GO:0051020 |
| 8 | 0.13 | 0.3425 | 2.46 | 0.00 | 0.73 | 3c4iA | GO:0003677 |
| 9 | 0.13 | 0.4595 | 2.43 | 0.08 | 1.00 | 1ashA | GO:0005506 GO:0015671 GO:0019825 GO:0020037 |
| 10 | 0.13 | 0.4699 | 2.11 | 0.20 | 0.92 | 3g4dA | GO:0008152 GO:0016829 GO:0000287 GO:0047461 GO:0046872 |

##### Consensus prediction of GO terms

**Molecular Function** GO:0003723 GO:0035639 GO:0016462 GO:0032561 GO:0046906 GO:0043167

| | | | | | | |
|---|---|---|---|---|---|---|
| **GO-Score** | 0.40 | 0.34 | 0.34 | 0.34 | 0.30 | 0.30 |
| **Biological Process** | GO:0006613 | GO:0045047 | GO:0015669 | | | |
| **GO-Score** | 0.34 | 0.34 | 0.30 | | | |
| **Cellular Component** | GO:0048500 | GO:0044445 | | | | |
| **GO-Score** | 0.34 | 0.30 | | | | |

(a) Cscore$^{GO}$ is a combined measure for evaluating global and local similarity between query and template protein. It's range is [0-1] and higher values indicate more confident predictions.

(b) TM-score is a measure of global structural similarity between query and template protein.

(c) RMSD$^{a}$ is the RMSD between residues that are structurally aligned by TM-align.

(d) IDEN$^{a}$ is the percentage sequence identity in the structurally aligned region.

(e) Cov represents the coverage of global structural alignment and is equal to the number of structurally aligned residues divided by length of the query protein.

(f) The second table shows a consensus GO terms amongst the top scoring templates. The GO-Score associated with each prediction is defined as the average weight of the GO term, where the weights are assigned based on Cscore$^{GO}$ of the template.

**[Click on S609161_results.tar.bz2 to download the tarball file including all modeling results listed on this page]**

---

Please cite the following articles when you use the I-TASSER server:

1. J Yang, Y Zhang. I-TASSER server: new development for protein structure and function predictions, Nucleic Acids Research, 43: W174-W181, 2015.
2. C Zhang, PL Freddolino, Y Zhang. COFACTOR: improved protein function prediction by combining structure, sequence and protein–protein interaction information. Nucleic Acids Research, 45: W291-W299, 2017.

Fig. S5

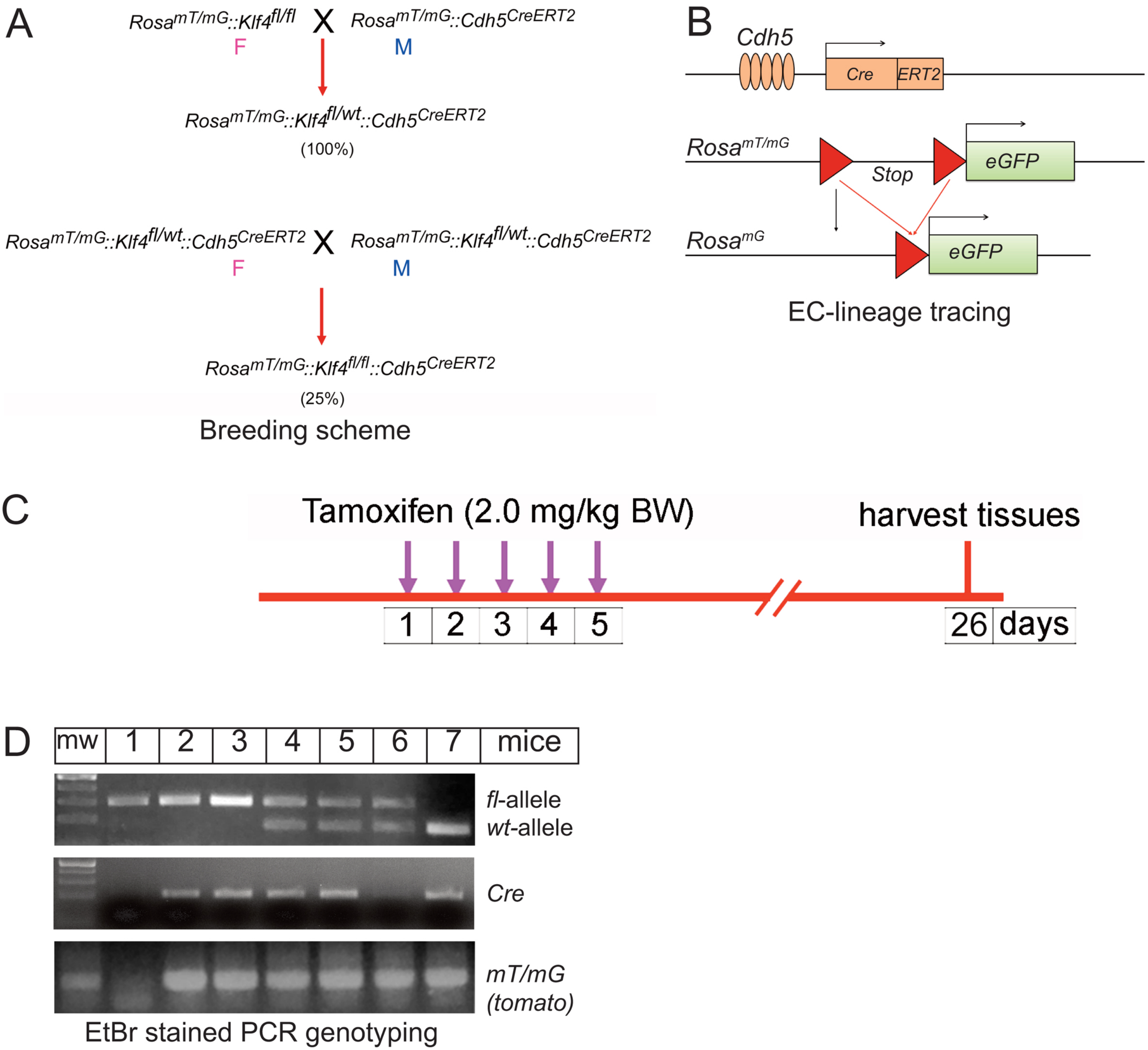
A
Rosa mT/mG::Klf4 fl/fl
X
Rosa mT/mG::Cdh5 CreERT2
F
M
Rosa mT/mG::Klf4 fl/wt::Cdh5 CreERT2
(100%)
Rosa mT/mG::Klf4 fl/wt::Cdh5 CreERT2
X
Rosa mT/mG::Klf4 fl/wt::Cdh5 CreERT2
F
M
Rosa mT/mG::Klf4 fl/fl::Cdh5 CreERT2
(25%)
Breeding scheme
B
Cdh5
Cre
ERT2
Rosa mT/mG
Stop
eGFP
Rosa mG
eGFP
EC-lineage tracing
C
Tamoxifen (2.0 mg/kg BW)
harvest tissues
1
2
3
4
5
26
days
D
mw
1
2
3
4
5
6
7
mice
fl-allele
wt-allele
Cre
mT/mG
(tomato)
EtBr stained PCR genotyping

**Table I**: Oligonucleotide primers used for PCR genotyping

| | **Primer names** | **Sequence (5'-3')** |
|---|---|---|
| 1 | Rosa$^{mT/mG}$ (TdTomato/Red) | *(F) 5'-AGCAAGGGCGAGGAGGTATC-3'*<br>*(R) 5'-CCTTGGAGCCGTACATGAACTGG-3'* |
| 2 | Cdh5$^{CreERT2}$ | *(F) 5'-GCGGTCTGGCAGTAAAAACTATC-3'*<br>*(R) 5'-GTGAAACAGCATTGCTGTCATT-3'* |
| | ***Mouse Klf4* specific primers** | **Sequence (5'-3')** |
| 3 | Klf4 exon 1 | *(F) 5'-CTGGGCCCCCACATTAATGAG-3'* |
| 4 | Klf4 exon 2 | *(R) 5'-CGCTGACAGCCATGTCAGACT-3'* |

Diagnostic genotyping PCR product sizes:

| | **Genotype** | **Amplicon size (bp)** |
|---|---|---|
| 1 | Wild type (wt) | 172 bp |
| 2 | Homozygote (hom) | 296 bp |
| 3 | Heterozygote (het) | 296 bp and 172 bp |

**Table 2**: **Primer Sequences for qRT-PCR**

| | **Name** | **Species** | **Sequence (5'-3')** | **Accession#** |
|---|---|---|---|---|
| **1** | *18S rRNA* | mouse | ***(F)*** *5'-GCAATTATTCCCCATGAACG-3'*<br>***(R)*** *5'-GGCCTCACTAAACCATCCAA-3'* | NR_003278 |
| **2** | *Klf4* | Mouse | ***(F)*** *5'-GAGGAAGCGATTCAGGTACAGAAC-3'*<br>***(R)*** *5'-AGGCTTATTTACCTGGCTTAGGTC-3'* | NM_010637.3 |
| **3** | *Klf2* | mouse | ***(F)*** *5'-CCTCCCAAACTGTGACTGGTAT-3'*<br>***(R)*** *5'-ATGCCCACCTGTCTCTCTATGT-3'* | NM_008452.2 |
| **4** | *Flk1/Vegfr2* | mouse | ***(F)*** *5'-TGTGGTCTCACTACCAGTTAAAGC-3'*<br>***(R)*** *5'-CATTCGATCCAGTTTCACAGAG-3'* | NM_010612.2 |
| **5** | *Cdh5 (VE-cadherin)* | mouse | ***(F)*** *5'- AGATCCCAGAAGAGCTAAGAGGAC-3'*<br>*(**R**) 5'- AGAAAAGGAAGAGTGAGTGACCAG-3'* | NM_009868.4 |
| **6** | *Vcam-1* | mouse | ***(F)*** *5'-CCCAGGTGGAGGTCTACTCA-3'*<br>***(R)*** *5'-CAGGATTTTGGGAGCTGGTA-3'* | X67783.1 |
| **7** | *CD31/Pecam-1* | mouse | ***(F)*** *5'-GAGACTCAGAGGCGCTAGTTAAT-3'*<br>***(F)*** *5'-CTAACCCAGTGATTGACAACAGA-3'* | NM_008816.2 |
| **8** | *eNos* | mouse | ***(F)*** *5'-AGACCTCCTGAGGACAGAGCTA-3'*<br>***(R)*** *5'-GAAAAGCTCTGGGTGCGTA-3'* | NM_008713.4 |
| **9** | *αSMA* | mouse | ***(F)*** *5'-CCCCTGAAGAGCATCGGACA-3'*<br>*(**R**) 5'-TGGCGGGGACATTGAAGGT-3'* | NM_007392.3 |
| **10** | *Tagln (sm22)* | mouse | ***(F)*** *5'-ATTGGTGAACAGCCTGTATCCT-3'*<br>***(R)*** *5'-CTCCACGGTAGTTTCCATCG-3'* | NM_011526.5 |
| **11** | *Erg (Ets transcription factor)* | mouse | ***(F)*** *5'- ATCCTGGGACCGACCAGTAG-3'*<br>***(R)*** *5'- GTGATGCAGTTGGAGTTGGAG-3'* | NM_001302152.1 |
| **12** | *Ace2* | mouse | ***(F)*** *5'-CTACAGGCCCTTCAGCAAAG-3'*<br>*(**R**) 5'-TGCCCAGACCCTAGAGTTGT-3'* | AB053182.1 |
| **13** | *Thrombomodulin (Thbd)* | mouse | *(**F**) 5'-TAGGGAAGACACCAAGGAAGAG-3'*<br>*(**R**) 5'-CTTGCGCAGGTGACAGAGAAG-3'* | NM_009378.3 |
| **14** | *Endothelin-1/Edn1* | mouse | ***(F)*** *5'-CCAAAGTACCATGCAGAAAAGC-3'*<br>***(R)*** *5'-CAGCTTTCAACTTTGCAACACG-3'* | NM_010104.4 |
| **15** | *IL-6* | mouse | ***(F)*** *5'-TACCACTTCACAAGTCGGAGGC-3'*<br>***(R)*** *5'-CTGCAAGTGCATCATCGTTGTTC-3'* | NM_001314054 |
| **16** | *β-Cat (Ctnnb1)* | mouse | ***(F)*** *5'-TGACACCTCCCAAGTCCTTT-3'*<br>***(R)*** *5'-TTGCATACTGCCCGTCAAT-3'* | NM_007614.3 |

**F**: forward primer; **R**: reverse primer;

**KEY RESOURCES TABLE**

| REAGENT OR RESOURCE | | | CATALOG# Or RRID# |
|---|---|---|---|
| **Antibodies** | **Dilution** | **SOURCE** | |
| Goat anti-mouse/rat CD31/PECAM-1 | 1:25 (staining) | R&D Systems | AF3628 |
| Goat anti-mouse/rat CD31/PECAM-1 | 1:100 (flow) | R&D Systems | AF3628 |
| Mouse monoclonal anti-KLF4 | 2 μg/ml (WB) | Rockland | 200-301-CE9 |
| Rabbit Lamin-B1 | 2 μg/ml (WB) | Rockland | 600-401-P62 |
| Mouse anti-α-smooth muscle actin | 2 μg/ml (WB) | Sigma-Aldrich | A2547 |
| Rat ant-CD31 (MEC7.46) | 1:25 (staining) | Abcam | Ab7388 |
| Rat ant-CD31 (MEC7.46) | 1:200 (flow) | Abcam | Ab7388 |
| Rabbit polyclonal anti-TAGLN/Transgelin | 1:25 (staining) | Abcam | ab14106 |
| Sheep anti-von Willebrand Factor (vWF) | 1:25 (staining) | Abcam | ab11713 |
| Rabbit anti-Collagen I | 2 mg/ml (WB) | Abcam | ab34710 |
| Anti-sheep IgG-Alexa Fluor 488 | 1:200 | Abcam | ab150177 |
| Anti-rabbit IgG-Alexa Fluor 647 | 1:200 | Abcam | ab150063 |
| Mouse monoclonal anti-Klf4 (F8) | 2 μg/ml (WB) | Santa Cruz | 166238 |
| Rabbit anti-KLF4 (H-180) | 2 μg/ml (WB) | Santa Cruz | sc-20691 |
| Rabbit anti-KLF2 (H-60) | 2 μg/ml (WB) | Santa Cruz | sc-28675 |
| Mouse monoclonal anti-TGF-β1 | 2 μg/ml (WB) | Santa Cruz | sc130348 |
| Mouse monoclonal anti-p300 (NM11) | 2 μg/ml (WB) | Santa Cruz | sc-32244 |
| Mouse anti-VEGFR2/FLK1 (A3) | 2 μg/ml (WB) | Santa Cruz | sc-6251 |
| Mouse monoclonal anti-Endoglin | 1:25 (staining) | Santa Cruz | sc-376381 |
| Monoclonal anti-Endoglin (P3D1) | 2 μg/ml (WB) | Santa Cruz | sc-18838 |
| Mouse anti-VE-cadherin (Cdh5/CD144) | 1 μg/ml (WB) | Santa Cruz | sc-9989 |
| Rat anti-mouse CD31 | 1:200 (flow) | BD Biosciences | 550274 |
| Rabbit anti-von Willebrand factor (vWF) | 1:40 (staining) | EMD Millipore | AB7356 |
| Rabbit anti-GAPDH | 1 μg/ml (WB) | Cell Signaling | 5174 |
| Rabbit anti-VEGFR2/FLK1 (55B11) | 2 μg/ml (WB) | Cell Signaling | 2479 |
| Rabbit anti-eNOS (D9A5L) | 2 μg/ml (WB) | Cell Signaling | 32027 |
| Rabbit anti-α-SMA (D4K9N) | 2 μg/ml (WB) | Cell Signaling | 19245S |
| Rabbit anti-P-Smad2 (S465/467) | 2 μg/ml (WB) | Cell Signaling | 3108S |
| Rabbit anti-mouse VCAM-1 (D8U5V) | 2 μg/ml (WB) | Cell Signaling | 39036 |
| Rabbit anti-human VCAM-1 (E1E8X) | 2 μg/ml (WB) | Cell Signaling | 13662 |
| Rabbit anti-FLAG (M2) (D6W5B) | 2 μg/ml (WB) | Cell Signaling | 14793 |
| Rabbit anti-FLAG (M2) (D6W5B) | 2 μg/IP (co-IP) | Cell Signaling | 14793 |
| Rat anti-mouse VE-cadherin (Cdh5/CD144) | 2 μg/ml (WB) | Thermo-Fisher | 14-1441-82 |
| Rat anti-mouse VE-cadherin (Cdh5/CD144) | 1:100 (flow) | Thermo-Fisher | 14-1441-82 |
| Anti-KLF4 antibody (11880-1-AP) | 2 μg/ml (WB) | Thermo-Fisher | PA5-27440 |
| Anti-KLF2 antibody | 2 μg/ml (WB) | Thermo-Fisher | PA5-40591 |
| Anti-ACE2 Monoclonal Antibody (CL4035) | 2 μg/ml (WB) | Thermo-Fisher | MA5-31395 |
| Rabbit anti-KLF4 | 3 μg/ml | Thermo-Fisher | PA5-35303 |
| Donkey anti-rabbit Alexa Flour-594 | 1:200 | Thermo-Fisher | AB-141637 |
| Chicken anti-rabbit Alexa Fluor-647 | 1:200 | Thermo-Fisher | AB-2535861 |
| Goat anti-mouse Alexa Flour-488 | 1:200 | Thermo-Fisher | AB-2534069 |

| | | | |
|---|---|---|---|
| Mouse anti-human KLF4 antibody | 2μg/IP | Novus | H00009314-M01 |
| Anti-rabbit IgG-TRITC | 1:200 | Novus | NBP1-75270 |
| Anti-mouse IgG-DyLight 594 | 1:200 | Novus | NBP1-75563 |
| Mouse anti-Tubulin (AA10) | 1 μg/ml (WB) | BioLegend | 657402 |
| Anti-mouse CD11b (clone M1/70) | 1:200 (flow) | BioLegend | 101206 |
| Anti-LY-6G (clone 1A8) | 1:200 (flow) | BioLegend | 127610 |
| Mouse anti-GST antibody (P1A12) | 1 μg/ml (WB) | BioLegend | 640802 |
| **Mice and Animals** | **Sex** | **SUPPLIER** | **STOCK#** |
| *Klf4*$^{wt/fl}$ | Breeder pair | MMRRC | 29877 |
| *Gt(Rosa)*$^{26Sortm4(ACTB-tdTomato,-EGFP)Luo/J}$ | Breeder pair | JaxLab | 007576 |
| *tg.Cdh5(PAC)*$^{CreERT2}$ | Breeder pair | UK Cancer Res | --- |
| **Cells** | **Abbreviation** | **SUPPLIER** | **CAT#** |
| Human umbilical vein endothelial cells | HUVECs | LONZA | C2519A |
| Human lung microvessel endothelial cell | hLMVECs | PromoCell | C-12281 |
| RAW 264.7 macrophage cells | Macrophages | Kostandin Pajcini | gift |
| HEK293T Packaging cell line | Phenix-Ampho | Genscript | T98134 |
| **Chemicals, Media and Reagents** | | **SUPPLIER** | **CAT#** |
| Lenti-X Concentrator | | Clontech/TAKARA | 631321 |
| Polybrene infection reagent | Polybrene | Merck | TR-1003-G |
| Tamoxifen | TAM | Sigma/Millipore | T5648 |
| Corn oil | | Sigma/Millipore | C8267 |
| Dulbecco's Modified Eagle Medium | DMEM | Sigma/Millipore | FG4815-BC |
| Endothelial Cell Media kit | EndoGRO | Sigma/Millipore | SCME004 |
| Elastase assay kit | | Sigma/Millipore | MAK246-1KT |
| Fetal Bovine Serum | FBS | Sigma/Millipore | TMS-013-B |
| L-Glutamine | L-Glu | Sigma/Millipore | K0282-BC |
| Antibiotic antimycotic solution (100x) | PenStrep | Sigma/Millipore | A5955-100ML |
| Gelatin from porcine skin | Gelatin | Sigma/Millipore | G2500-100G |
| Bis-Acrylamide solution (1:30) | PAGE | BioRad | 1610156 |
| Tetramethylethylenediamine | TEMED | BioRad | 161-0801 |
| Alpha-1-Anti-Trypsin (A1AT) | Prolastin | Grifolis | -- |
| Microbeads coupled mouse CD45 antibody | CD45-beads | Miltenyi Biotech | 130-052-301 |
| Microbeads coupled mouse CD31 antibody | CD31-beads | Miltenyi Biotech | 130-097-418 |
| Secrete-Pair kit | | GeneCopoeia | LF031 |
| SYBR Green master mix | | Applied Biosystm | 4385612 |
| Apoptosis-detection kit APC | | eBiosciences | 88-8007-72 |
| ELISA kit, IL-1β | | ThermoFisher | BMS6002 |
| ELISA kit, TNFα | | ThermoFisher | BMS607-3 |
| ELISA kit, IL-6 | | ThermoFisher | KMC0061 |
| Puromycin | 6μg/ml | ThermoFisher | A11138-03 |
| Prolonged Gold DAPI with antifade | | ThermoFisher | P36962 |

| **Plasmids and vectors** | | **SUPPLIER** | **CAT#** |
|---|---|---|---|
| pLNCX2 (lentivirus) | DNAs | Takara | 631503 |
| Klf4-shRNA | Lentivirus | OpenBiosystems | --- |
| pLenti-C-Myc-DDK (PS100064) | Lentivirus | Origin | --- |
| Human KLF4 cDNA (NM_004235) | Lentivirus | Origin | RC206691L1 |
| Human KLF2 cDNA ((NM_016270) | Lentivirus | Origin | RC210042L1 |
| pEZX-LvG04 | Virus vector | GeneCopoeia | custom |
| pLNCX2, pLNCX2-shRNAs | DNAs | Genscript | custom |
| KLF4-WT and mutant-deletion constructs | DNAs | Genscript | custom |
| GST-KLF4 fusion proteins constructs | DNAs | Genscript | custom |
